# Cross-modal influence of odorant stimulation on sound localization revealed by EEG and diffusion modeling

**DOI:** 10.1101/2024.03.27.586970

**Authors:** Laura-Isabelle Klatt, Christine Ida Hucke

## Abstract

The ventriloquism effect, where sounds are mislocalized towards a spatially disparate but temporally synchronous visual stimulus, is well established. Liang et al. (2022) provided first evidence that chemosensory cues can similarly bias sound localization. This study, conducted in 2023, replicates and extends their behavioral findings, using diffusion modeling and EEG to identify underling computational and neural mechanisms. Participants localized (left vs. right) each sound in eight-item sequences, with trigeminally-potent odorants presented concurrently with the first four sounds (left, right, or both nostrils). Unbeknownst to participants, some sounds originated from the center. For central sounds, the proportion of right-ward responses increased with right-nostril stimulation but decreased with left-nostril stimulation (relative to controls). This bias was strongest in the second half of the sequence and disappeared at larger sound eccentricity. Diffusion modeling showed the bias was best explained by changes in drift rate (perceptual evidence accumulation), not in starting point (decision bias). EEG alpha-band activity lateralized towards the chosen side for central sounds, consistent with attentional orienting towards the perceived location, and was diminished at large eccentricities with incongruent chemosensory stimulation. The findings corroborate an odorant-sound spatial ventriloquism effect and provide novel insights into the cognitive and attentional processes involved.

**Public significance statement:** Our findings reveal that trigeminally potent odors can subtly influence sound localization judgments, even when those odors are irrelevant to the task at hand. This influence is strongest when sound information is uncertain (i.e., when unbeknownst to participants, a sound that had to be localized as coming from the left or right side, originated from a central location), suggesting that trigeminal stimulation can act as a spatial cue under ambiguous conditions. In addition, concurrent EEG cording revealed that the influence of odorant stimulation was also reflected in neural correlates of auditory spatial attention. Critically, diffusion modeling showed that the observed cross-modal bias occurs at a perceptual level. These findings highlight that the human brain integrates information from the chemical sense and hearing more deeply than previously assumed, revealing that even task-irrelevant odors can shape spatial perception in other senses.

---

Although humans typically prioritize vision or audition to extract spatial information from the environment, the chemical sense – including olfaction and trigeminal sensations – can also provide spatial information. For instance, chemical cues can be utilized during wayfinding or object localization(Raithel & Gottfried, 2021a; Welge-Lüssen et al., 2014; Wu et al., 2020) and are critical in identifying potential threats or resources, in particular when the availability of visual or auditory cues may be limited or ambiguous. Therefore, understanding how spatial information originating from the chemical senses is processed and integrated with other sensory inputs is vital to comprehend human multisensory perception.

Two primary mechanisms underlie the extraction of chemical spatial information from airborne molecules. Firstly, spatial information can be derived from serial comparisons of odorant concentration or intensity across consecutive sniffs while moving in space (Jacobs et al., 2015). Secondly, spatial localization while remaining stable can arise through intensity comparisons across nostrils or via monorhinal perception(Raithel & Gottfried, 2021b).

When conflicting sensory information is processed through different senses, we frequently observe that one sensory modality influences another. For instance, in the audiovisual domain, the ‘spatial ventriloquism effect’ describes how the perceived location of a sound is shifted towards a concurrently presented but spatially disparate visual stimulus (for a review, see(Chen & Vroomen, 2013)). Similar effects have been demonstrated in other modalities, such as the visuo-tactile domain(Bruns & Röder, 2010). However, only a limited number of studies have examined cross-modal interactions between the chemical and other human senses.

For instance, it has been shown that odorant stimulation can modulate visual motion processing (Kuang & Zhang, 2014) as well as visuo-spatial navigation (Wu et al., 2020). In the auditory domain, a behavioral study by Liang and colleagues (2022) recently investigated whether odorants can bias the perceived location of a centrally presented sound towards the stimulated nostril. Specifically, while performing a two-alternative forced choice sound localization task (left versus right), the participants’ left or right nostril was stimulated with a pure olfactory stimulus (phenylethyl alcohol, rose smell) or a bimodal (i.e., olfactory-trigeminal) stimulus (menthol, mint smell). The authors found that centrally presented sounds were more frequently categorized as originating from the right side if participants received monorhinal rightward odorant stimulation. Critically, this odorant-induced bias was only present with a bimodal olfactory-trigeminal stimulus but not with a pure olfactory stimulus. This is consistent with the finding that humans can only consciously localize monorhinally presented odors when they stimulate the trigeminal system (Croy et al., 2014; Kleemann et al., 2009; Kobal et al., 1989). Thus, trigeminal stimulation seems essential to evoke a biased spatial perception or lateralization of other senses such as audition.

While the findings by Liang and colleagues (2022) clearly demonstrate the influence of trigeminal cues on auditory localization judgements, the underlying cognitive mechanisms remain ambiguous. Although the authors interpret their results as evidence of odor-sound integration, alternative explanations such as decision bias or attentional cueing have not been convincingly ruled out.

Hence, the current study has three main objectives: First, we aim to replicate the behavioral odorant-induced sound localization bias reported by Liang et al.(2022), using a different bimodal stimulus (i.e., isopropanol) and a modified, EEG-compatible design. Second, we apply drift diffusion modeling (Ratcliff & McKoon, 2008) to determine whether the observed bias reflects a decision-level shift (i.e., a change in starting point towards the stimulated nostril), enhanced evidence accumulation (i.e., increased drift rate), or both. Finally, Liang et al. (2022) argued against a role of selective attention in explaining their results based on the fact that both confidence ratings and sound localization ratings were in line with the principle of inverse effectiveness in multisensory integration. Here, we draw on electrophysiological data to examine the role of spatial attention more explicitly. We investigate whether chemosensory cues influence auditory spatial attention by analyzing hemispheric modulations of alpha power. Alpha power lateralization has been linked to the allocation of spatial attention in various domains, such as anticipatory shifts of spatial attention (Rihs et al., 2007; Sauseng et al., 2005; Thut et al., 2006; Yamagishi et al., 2005), attentional orienting within working memory (Klatt et al., 2020; Myers et al., 2015; Poch et al., 2017; Schneider et al., 2016), or attentional selection from a stimulus array (Bacigalupo & Luck, 2019; Getzmann et al., 2020).

In this study, participants completed a sound localization paradigm, in which they were presented with a sequence of broadband pink noise bursts from various lateralized locations (see Figure 1-B). After each sound presentation, participants made a two-alternative forced choice (2AFC) judgement, indicating the spatial location of the sound as coming from the left or right side. Critically, on a subset of occasions, but unbeknown to the participants, the sounds were presented at a central location, creating a situation of maximal uncertainty. Note that, absolute sound localization (i.e., indicating the apparent stimulus location via head movement or pointing) is known to be most precise for sounds presented in front of the subject with localization errors increasing for more peripheral stimulus locations (Makous & Middlebrooks, 1990). In contrast, the present study employed a relative sound localization judgement, in which participants typically exhibit reduced precision for locations close to midline (Golob et al., 2017; Liang et al., 2022). Accordingly, in the following, we refer to central sound locations as providing uncertain or ambiguous location information. The auditory stimuli were grouped into a sequence of eight consecutive sounds (Figure 1C). Concurrently, during the first half of the sequence, sound presentation could be accompanied by trigeminally potent odorant stimulation in the left, right, or both nostrils (Figure 1D). In addition to the birhinal stimulation condition, a subset of auditory-only trials served as a neutral control condition. Participants were informed that the chemosensory stimulation was task-irrelevant.

**Figure 1.**
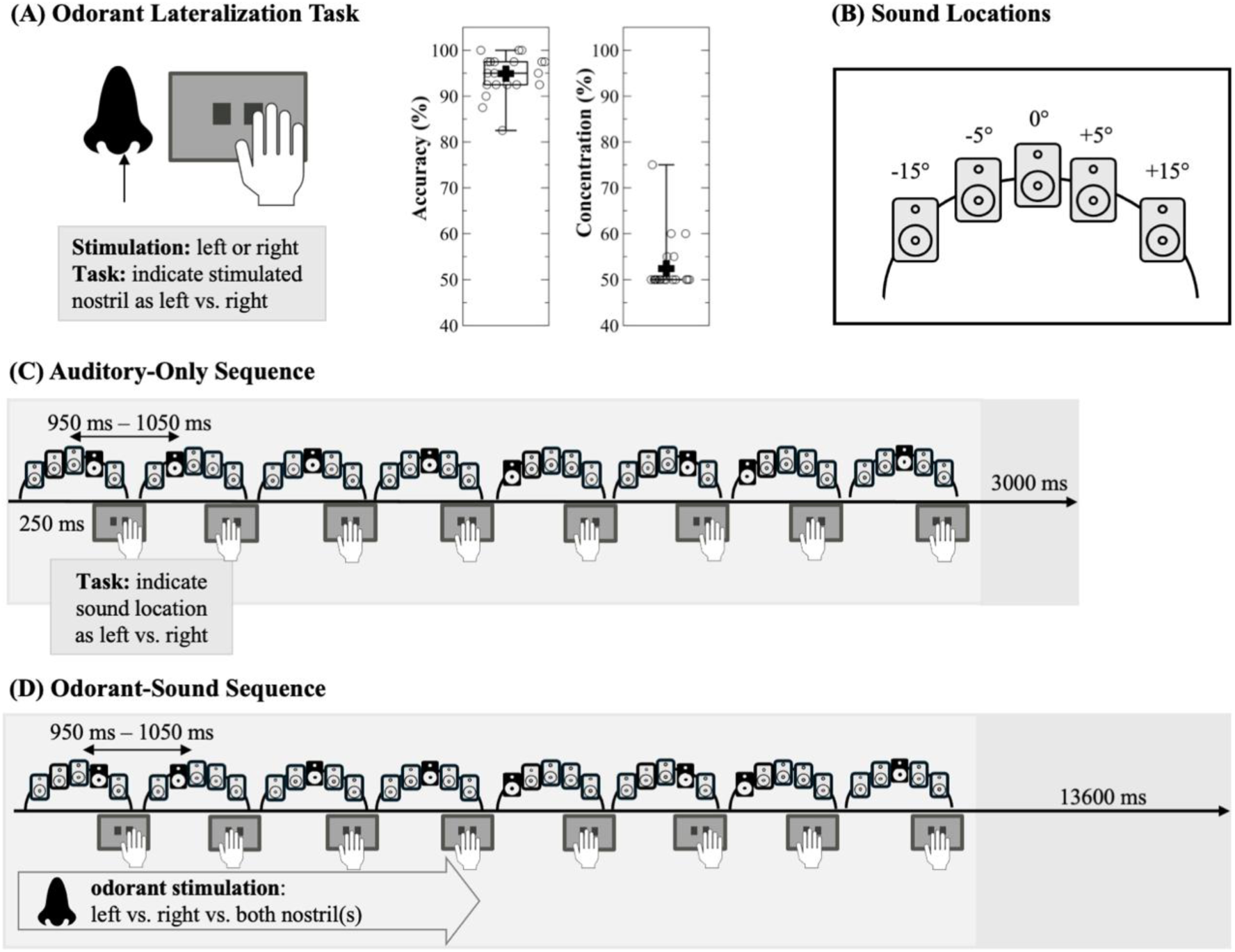
Experimental Design. (A) To determine the concentration that is trigeminally potent for each individual subject, participants completed an odorant lateralization task in the beginning of the experiment, in which they indicated whether the left or right nostril was stimulated. Boxplots show the distribution of task accuracy and final concentration thresholds, with horizontal lines marking the median, the plus indicating the group mean, and whiskers denoting +-1.5 IQR. Dots show individual participant mean accuracy or concentration threshold. (B) During the sound localization paradigm, sounds could be presented at five different locations (-15°, -5°, 0°, +5°, -15° azimuth in the horizontal plane). Speaker locations were covered by an opaque curtain to not reveal the presence of the central speaker to participants. (C) Schematic illustration of an auditory-only sound sequence, consisting of eight broadband pink noise sounds bursts. The activated loudspeaker is highlighted in black, while non-activated loudspeakers are displayed in grey. For each noise burst in a sequence, participants were required to indicate the perceived direction of the sound (left vs. right) via button press with their dominant hand. The inter stimulus interval was between 950 and 1050 ms and the inter sequence interval was 3 s. (D) Illustration of a sound sequence that was accompanied by a 4 s bimodal odorant stimulation either in the left, right, or both nostril(s) overlapping with the presentation of the first 4 sounds. The inter-sequence interval was 13.6 s.

In line with the results by Liang and colleagues (2022), we expect that monorhinal stimulation biases the reported sound localization towards the stimulated nostril, especially under conditions of high uncertainty (i.e. centrally presented sounds). If chemosensory stimulation leads to a decisional bias towards the stimulated nostril side, this should manifest itself in the starting point parameter of the diffusion model, whereas a perceptual bias should influence the drift rate (i.e., the rate of evidence accumulation). At the electrophysiological level, we test whether alpha power lateralization, an index of auditory spatial attention, is modulated by the spatial congruency between odorant and sound cues.

## Methods

### Power Analysis

We conducted an a-priori power analysis, using MorePower 6.0(Campbell & Thompson, 2012). Liang et al.(Liang et al., 2022) reported an effect of right-nostril menthol stimulation on the proportion of right-ward responses for sounds at azimuth = 0° compared to a control condition (*d* = 0.608, i.e., *η*_p_ ^2^ = 0.2752). Accordingly, aiming for a power of 80% and an effect size of *η*_p_ ^2^ = 0.275 in a paired-sample *t*-test, the power analysis yields a required sample size of *n* = 24.

### Participants

A total of 32 participants were recruited for this study. Data collection took place in 2023 at the Leibniz Research Centre for Working Environment and Human Factors in Dortmund, Germany. A subset of 8 participants were students in a practical EEG seminar, taught for the Ruhr University Bochum in the summer semester of 2023. Exclusion criteria entailed migraine, pregnancy, history of neurological or psychiatric diseases, asthma, acute or chronic upper airway diseases, acute allergies affecting the respiratory system, current influence of a drug, alcohol, or a controlled substance, impaired hearing ability or olfaction, as well as a hairstyle incompatible with EEG recordings. Hearing levels of all participants were assessed by means of a pure tone audiometry (Oscilla USB 330; Immedico, Lystrup, Denmark). The fully automated procedure included the presentation of eleven pure tones at varying frequencies (i.e., 125 Hz, 250 Hz, 500 Hz, 750 Hz, 1000 Hz, 1500 Hz, 2000 Hz, 3000 Hz, 4000 Hz, 6000 Hz, 8000 Hz). In addition, on the day of the experiment, participants were required to pass a pulmonary lung function following to the ATS/ERS 2019 spirometry standards (Graham et al., 2019), that is, forced exhaled volume (FEV1) must surpass 85% (Vyaire Sentry Suite). Moreover, an olfactory function test was conducted (Sniffin’ Sticks identification subtest, Burghart Messtechnik GmbH, Wedel, Germany; Hummel et al., 2007; Oleszkiewicz et al., 2019). In addition, the nasal flow rate of each nostril was examined by means of an active anterior rhinomanometry (RHINO-SYS, Happersberger otopront GmbH, Hohenstein, Germany) to determine if participants showed signs of a nasal flow asymmetry.

Three participants were excluded because the odorant lateralization paradigm (see section 2.4.1) had to be aborted prior to reaching the required performance level. Further, two participants were excluded due to insufficient lung function (spirometry, forced exhaled volume < 85%). In addition, one participant was not able to properly execute the velopharyngeal closure technique, while another participant was excluded for the intake of psychotropic drugs. That is, the final sample included 25 participants (male = 12, female = 13).

The mean age in the sample was 24.8 years (*SD* = 3.52). 23 participants reported to be right-handed, two participants were left-handed. The majority of participants (*n* = 20) indicated to be non-smokers, 4 participants indicated to smoke only occasionally, while 1 participant indicated that they smoked on a regular basis. Note that smoking did not result in exclusion, as long as participants passed the odorant lateralization paradigm, as described in section 2.4.1. Due to a technical error, audiometry data for two subjects were not saved. Considering all other subjects in the sample, hearing thresholds for all tested frequencies were ≤ 25 dB in 22 subjects, indicating unimpaired hearing (Olusanya et al., 2019). Four subjects showed negligible outliers at 30 dB (n = 3) and 35 dB (n = 1) for one of the tested frequencies, while all other frequencies fell below 30 dB. Since the stimuli used in the present study were broad-band (see section 2.3), those outliers were considered negligible. The average forced exhaled volume in the sample was 101.04% (*SD* = 10.19). On the Sniffin’ Sticks olfactory function test, participants scored on average 12.13 out of 16 points (*SD* = 1.72). The flow data of the two nostrils did not differ significantly, neither for the influx, *t*(22) = 1.26, *p* = .22, nor for the efflux, *t*(22) = 1.35, *p* = .19. Thus, there was no systematic bias with respect to the stimulated nostril.

All participants provided informed consent prior to the beginning of the experimental procedure. The study was approved by the local ethics committee at the Leibniz Research Centre for Working Environment and Human factors (Application No. 242) and conducted in accordance with the Declaration of Helsinki. As compensation for their time, participants received 12 Euros per hour (or course credit).

### Apparatus and Stimuli

The experiment took place in a dimly lit, sound attenuated room (5.0 × 3.3 × 2.4 m) with pyramid-shaped foam panels on ceiling and walls and a woolen carpet on the floor to dampen the background noise level (i.e., below 20 dB(A)). The experiment was programmed and controlled using ePrime 3.0 (Psychology Software Tools, Pittsburgh, PA) on a Windows 10 computer. The synchronization of auditory stimuli and EEG triggers was controlled using an AudioFile Stimulus Processor (Cambridge Research Systems, Rochester, UK). Participants were seated in a comfortable chair at a distance of approximately 130 cm from a 49’’ centrally aligned 1800R curved monitor (5120 × 1440 pixel resolution, 100 Hz refresh rate, Samsung, Seoul, South Korea), while placing their head on a chin rest that prevents head movements during the experiment. Below the screen, five full-range loudspeakers (SC 55.9 – 8 Ohm; Visaton, Haan, Germany) were mounted at -15°, -5°, 0°, +5°, and +15° horizontal azimuth (Figure 1B). Loudspeaker locations at -5° and +5° are also referred to as “intermediate positions”, whereas the loudspeaker locations at -15° and +15° are referred to as “outer positions”. To prevent participants from knowing the exact loudspeaker positions, the loudspeakers were covered by an opaque, sound-permeable curtain. Critically, the participants were not aware of the presence of the central loudspeaker. As auditory stimuli, a 250 ms broad-band pink noise was generated. All sound stimuli were presented at a sound level of ∼61 dB(A).

Isopropanol, dissolved in water at an individualized concentration (ranging from 50% - 75%) was used as a bimodal odorant substance. Isopropanol is a commercially available substance which is used amongst others in making cosmetics, skin or hair products, perfumes, and disinfectants. The trigeminal component of the bimodal odorant elicits an immediate short-lasting irritating sensation in most subjects (Smeets et al., 2002; Smeets & Dalton, 2002). For olfactory stimulus presentation, an EEG-compatible computer-controlled flow-olfactometer with a constant air flow of 2.5 l/min (NeuroDevice, Version 2, Warsaw, Poland) was used. For an elaborate description of the device and for technical adaptions made in order to optimize the setup for concurrent EEG recordings, please see Hucke et al (Hucke et al., 2018, 2021), respectively. Briefly, the olfactometer delivers olfactory stimuli into the participants nostrils via a nasal cannula. To allow for monorhinal odorant stimulation, a custom-built clip separates the two nasal tubes, however, birhinal stimulation is possible as well. The olfactometer integrated the odorous stimulus into the constant air flow of clean air and thereby ensures a seamless and precise stimulus on- and offset. This mechanism spares a potential co-activation of mechanoreceptors of the trigeminal nerve.

### Procedure and experimental protocols

Prior to the experiment, participants were informed about the experimental procedure and the potentially irritating effects of the stimulation with isopropanol. First, all participants were screened for exclusion and inclusion criteria and the required pre-assessments were conducted (see section 2.2). Afterwards, participants were prepared for the EEG recording and instructed to practice the velopharyngeal closure technique. The latter refers to a breathing technique that aims to seal the velum and pharyngeal walls and prevent air flow from the oral to the nasal cavities while breathing through the mouth(Kobal, 1981, p. 19). This ensures that olfactory stimulation is not affected by the participant’s respiration. To ensure trigeminal involvement, first, participants’ odorant lateralization threshold was determined (as described in section 2.4.1). Once the final concentration was determined, participants continued with the 2AFC sound localization task (as described in section 2.4.2). The entire experimental procedure (including EEG cap preparation) took approximately 2 - 2 ½ hours.

### Odorant lateralization paradigm

To ensure that all participants were able to correctly localize the bimodal olfactory stimulus, a lateralization paradigm was conducted as a confirmation of trigeminal involvement (Figure 1A). The latter comprised an iterative procedure in which participants were asked to indicate whether a monorhinally-presented isopropanol stimulus was perceived in the left vs. right nostril. All participants started with a concentration of 50% isopropanol. Odorant stimulation occurred randomly in the left or right nostril for 4 seconds and was followed by a stimulation-free inter-trial interval of 10 seconds (plus a random jitter of 0-100 ms). To make sure that participants’ attention was focused at the beginning of each trial, despite the long inter-trial interval, the fixation cross briefly increased in size 1000 ms prior to odorant stimulation onset to signal participants to anticipate the next odorant stimulus. If participants did not reach a lateralization accuracy of at least 80% in a block of 20 trials, the concentration was increased by 5%. To pass the lateralization test, accuracy had to be at least 80% on average in two consecutive blocks (40 trials) with the same isopropanol concentration(Mai et al., 2025). In the final sample, the median isopropanol concentration was 50% (range = 50-75%, cf. Figure 1A).

### Sound localization paradigm

A schematic sound sequence of the 2AFC auditory localization paradigm is depicted in Figure 1C and D. The auditory stimuli are grouped into sequences of 8 pink noise bursts with an inter-stimulus-interval of 950 ms (+ 0-100 ms random jitter). This amounts to roughly 8 s for a sound sequence. The order of sound locations within a sequence was counterbalanced and pseudo-randomized across all trials of the experiment and conditions (auditory-only, birhinal, odorant-left, odorant-right) such that 50% of all sound stimuli were presented at 0° azimuth and in 12.5% of cases at each lateralized position (+/- 15°, +/- 5°). This was motivated by the fact that the main (behavioral analysis) focused on the localization of sounds at the central position. For details on how the trials were sorted into the conditions, please refer to Table 1. In the analysis, we will further differentiate between sounds in the first (sounds 1-4) as opposed to the second half (sounds 5-8) of the sequence. The participants were asked to indicate the perceived direction (left vs. right) of each sound in a sequence. Participants responded as fast as possible via button press using the index and middle finger of the dominant hand. The response window ended with the onset of the next sound or after 950 ms in case of the last sound of the sequence. Participants were informed that some sounds would be easier, others very hard to categorize as coming from the left or right side to prevent frustration due to the fact that centrally presented sounds did not fall in either category. A total of eight sequences constituted one block. Each block was assigned to one of the following conditions: (i) auditory-only, (ii) birhinal, (iii) monorhinally left, or (iv) right isopropanol stimulation. Each odorant sequence contained a respective odorant stimulation for 4 s that was time-locked to the first sound in the sequence (Figure 1D). Due to the rising time of the olfactometer, the molecules build-up in the nasal mucus, and the overall slower and more long-lasting chemosensory processing it was expected that the odorant perception persists after stimulation offset and until the end of the sound sequence(Carlson et al., 2013; Hummel et al., 1992). In blocks including odorant stimulation, the inter-sequence-interval was 13.6 s to minimize habituation or sensitization. In auditory-only blocks, the inter-sequence-interval was 3 s. 1000 ms prior to the first stimulus presentation in a sequence, the fixation briefly increased in size to signal the beginning of the next sound sequence. Participants completed a total of 8 auditory-only blocks, and 12 odor-blocks in a randomized order. After 4 blocks, participants were invited to take a self-paced break. Participants were presented with 5 practice auditory-only sequences to familiarize themselves with the sequence structure and pace. Participants were instructed to breathe through their mouth while performing the velopharyngeal closure technique throughout the experiment. To dampen discomfort due to dry mouth feel, participants were encouraged to drink water and switch to nasal breathing during the breaks.

**Table 1.**
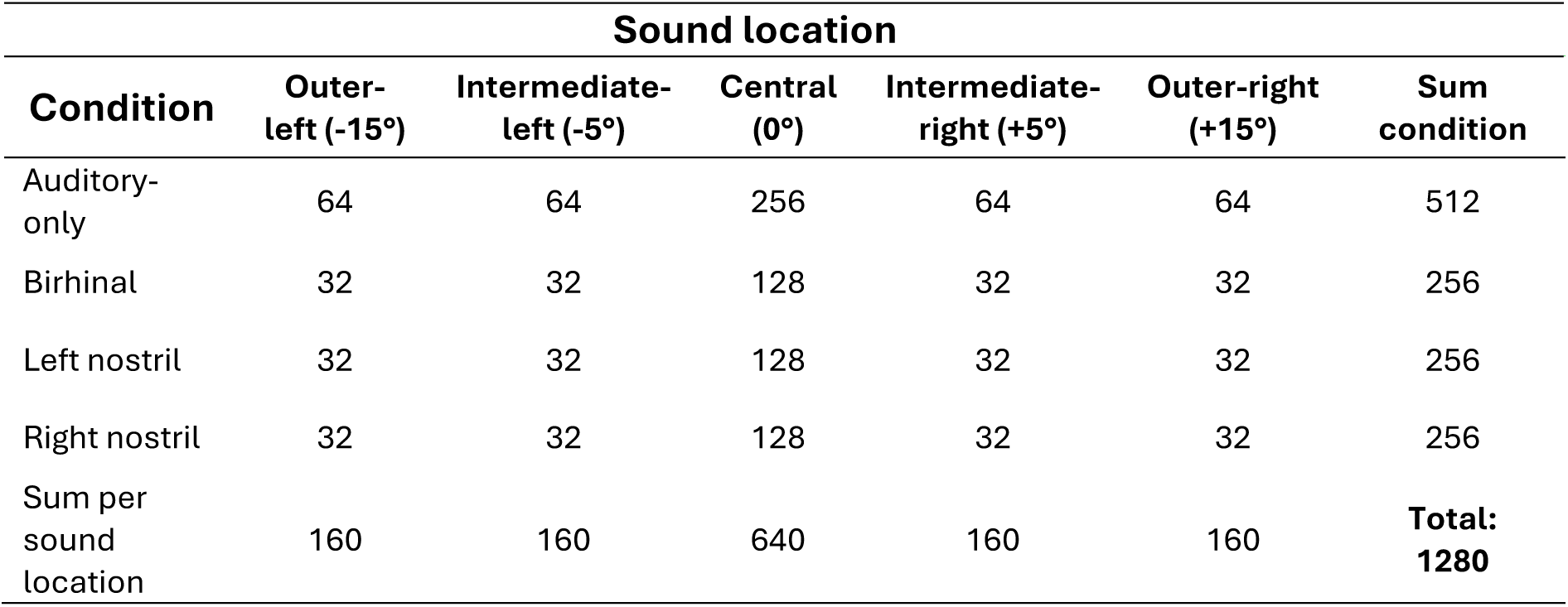
Trial numbers split according to the sound locations (5 speakers) and stimulation (auditory-only, birhinal, left-nostril, right-nostril) and the respective sums across the locations and conditions.

| Condition | Sound location |  |  |  |  | Sum condition |
| --- | --- | --- | --- | --- | --- | --- |
|  | Outer-left (-15°) | Intermediate-left (-5°) | Central (0°) | Intermediate-right (+5°) | Outer-right (+15°) |  |
| Auditory-only | 64 | 64 | 256 | 64 | 64 | 512 |
| Birhinal | 32 | 32 | 128 | 32 | 32 | 256 |
| Left nostril | 32 | 32 | 128 | 32 | 32 | 256 |
| Right nostril | 32 | 32 | 128 | 32 | 32 | 256 |
| Sum per sound location | 160 | 160 | 640 | 160 | 160 | **Total: 1280** |

### EEG data acquisition

The electroencephalogram was recorded using 64 Ag/AgCl electrodes (BrainCap, Brainvision, Gilching, Germany) and digitized with a 1000 Hz sampling rate, using a NeurOne Tesla amplifier (Bittium Biosignals Ltd, Kuopio, Finland). The electrode arrangement across the scalp was in accordance with the extended international 10-20 system. AFz and FCz served as the online ground and reference electrode, respectively. During cap preparation, electrode impedances below 20 kΩ were ensured.

### EEG preprocessing

The raw EEG data was preprocessed and cleaned from artifacts using a customized, automated preprocessing pipeline, implemented in MATLAB (R2021b and R2023a) and EEGLAB (2021.1; Delorme & Makeig, 2004). First, a non-causal, zero-phase Hamming windowed sinc FIR high-pass (pass-band edge: 0.01 Hz, transition band width: 0.01 Hz, filter order: 330000, -6dB cutoff frequency: 0.005 Hz) and low-pass filter (pass-band edge: 40 Hz, transition band width: 10 Hz, filter order: 33, -6dB cutoff frequency: 45 Hz) were applied to the continuous data. Then, electrodes compromised by artifacts were identified and excluded based on statistical properties of the data (i.e., normalized kurtosis value exceeding 5 standard deviations of the mean. In addition, any electrodes with periods of flatline recordings of more than 5 seconds were removed. On average, 4.48 electrodes were removed per subject (*SD* = 1.69). To restore the data at previously removed electrodes, a spherical interpolation algorithm was applied. In a next step, the data was re-referenced to the average of all channels, while retaining the original reference channel FCz in the dataset. At this point, a copy of the dataset was created and optimized for an independent component analysis (ICA)-based artefact identification procedure. Specifically, prior to ICA, the data was down sampled to 200 Hz and high-pass filtered at 1 Hz (pass-band edge, transition band width: 1 Hz, filter order: 3300, -6dB cutoff frequency: 0.5 Hz). The latter has been shown to increase the percentage of ‘near-dipolar’ independent components (ICs; (Winkler et al., 2015)). Then, the data was segmented into epochs ranging from -1500 ms to 1500 ms relative to each sound onset and submitted to an automated trial-rejection procedure. At this stage, on average 122 trials were rejected per subject (*SD* = 83.73). The remaining trials were submitted to a rank-reduced ICA. That is, by decomposing a principal component subspace of the data, the number of components to be decomposed was reduced by the number of rejected channels +1. This accounts for the rank deficiency in the data caused by the interpolation of rejected channels and the average reference procedure. To differentiate independent components (ICs) reflecting brain sources from those reflecting non-brain sources, the automated classifier tool ICLabel (Pion-Tonachini et al., 2019) was applied. For each IC, ICLabel assigned a probability value to each of following classes: “brain”, “eye”, “muscle”, “line noise”, “channel noise”, and “other”. Components with a probability estimate of > 30% in the category ‘eye’ or < 30% in the category ‘brain’ were flagged for rejection. However, prior to removal, the obtained ICA decomposition was copied to the unaltered reference dataset obtained prior to ICA-based artefact identification (i.e., sampling rate of 1000 Hz and high-pass filtered at 0.01 Hz). This dataset was segmented into epochs, as described above. Then, the ICs previously flagged as “artifactual” were subtracted. On average, 31.36 ICs were removed per subject (*SD* = 6.02). In a final step, trials that still contained large voltage fluctuations (±150 µV) were removed from the data. At this stage, on average 9 trials were rejected per subject (*SD* = 12.00).

### Time-Frequency Decomposition

The time-frequency decomposition was obtained using the EEGLAB built-in STUDY functions, applying Morlet Wavelet Convolution. Specifically, the preprocessed single-trial EEG data was convolved with a series of complex Morlet wavelets, varying in frequency from 4 to 30 Hz in 52 logarithmically spaced steps. A complex Morlet wavelet can be described as a Gaussian-modulated complex sine wave, where the number of cycles determines the width of the tapering Gaussian. To account for the trade-off between temporal and frequency precision, the number of cycles increased linearly as a function of frequency by a factor of 0.5, starting from 3 cycles at the lowest frequency (4 Hz) and up to 11.25 cycles at the highest frequency (30 Hz). The resulting event-related spectral perturbations (ERSPs) ranged from -1082 to 1082 ms relative to sound onset. We did not subtract a spectral baseline.

### Analysis

For repeated-measures ANOVA (rmANOVA), we report partial eta squared (*η*_p_ ^2^) as a measure of effect size. Mauchly’s test was used to assess the assumption of sphericity; in case of a violation (*p* < .05), Greenhouse-Geisser correction was applied. For directional a-priori hypotheses, that directly replicate previous tests by Liang et al(Liang et al., 2022), planned contrasts (using the ANOVA’s pooled error term) were computed. As JASP does not offer one-sided contrasts, the *p*-value was divided by two if the difference in means was in line with the direction of our hypothesis. For all other pairwise comparisons, one-sided paired sample *t*-tests were conducted and corrected for multiple comparisons using false discovery rate correction(Benjamini & Hochberg, 1995). Corrected *p*-values are denoted as *p*_corr_. Cohen’s *d_z_* values, using the standard deviation of the difference score as a denominator, are reported for both planned contrasts and paired sample *t*-tests. The alpha level for all tests was 0.05.

### Behavioral Performance

#### Central Sounds

For central sounds, participants were required to make a forced choice categorizing each sound as coming from the left or right side. Accordingly, there are no correct responses. Instead, the proportion of right-ward judgements served as a dependent variable to assess the influence of chemosensory stimulations on the response choice. Specifically, a 2-way rmANOVA with the within-subject factors *stimulation* (left nostril, right nostril, birhinal, auditory-only) and *sequence half* (first, second) was conducted.

In addition, reaction times to central sound localizations were analyzed using a 2-way rmANOVA with factors *sequence half* (first, second) and *stimulation* (congruent, incongruent, birhinal, auditory-only). Odorant stimulation could be either congruent with the given response (i.e., left nostril stimulation and left-ward response; right-nostril stimulation and right-ward response) or incongruent (i.e., left-nostril stimulation and right-ward responses; right-nostril stimulation and left-ward response). For the two control conditions (birhinal, auditory-only), response times were averaged across left and right button presses.

In addition, for both sets of analyses, planned one-sided contrasts were conducted to test our directional hypotheses based on results by Liang et al.(Liang et al., 2022). Specifically, we expected an increase of right-ward judgements during right-nostril stimulation and a decrease of right-ward indications during left-nostril stimulation. Analogously, faster responses are expected when participants classify a centrally presented sound in accordance with the side of odorant stimulation (congruent trials) as opposed to incongruent responses or responses in the control conditions. Thus, conditions with monorhinal stimulation were contrasted with the two control conditions.

#### Lateral Sounds

The influence of chemosensory stimulation on the correct categorization of lateral sounds was analyzed using a 3-way rmANOVA with the within-subject factors *stimulation* (congruent, incongruent, birhinal, auditory-only), *sound positions* (±15° azimuth, ± 5° azimuth) and *sequence halves* (first, second). Note that congruent and incongruent stimulation conditions refer to the side of the chemosensory stimulation relative to the actual sound location. That is, congruent trials include left-nostril stimulation and left responses as well as right-nostril stimulation and right responses, whereas incongruent trials include left-nostril stimulation and right responses as well as right-nostril stimulation and left-response.

The influence of chemosensory stimulation on reaction times (RTs) to lateral sounds were analogously analyzed to the accuracy analysis by means of a 3-way rmANOVA. Only correct responses were considered in this analysis.

Again, for both analyses, planned one-sided contrasts were conducted guided by our hypotheses based on results by Liang et al.(Liang et al., 2022) The impact of odorant stimulation on sound localization was expected to substantial decrease or potentially disappear at farther lateralized sound locations with little to no uncertainty.

#### Control analyses

A subset of participants exhibited a strong bias towards one response side, even in the absence of any spatial cues (i.e., in auditory-only trials). To examine if the behavioral results were influenced by the strong decision tendency in part of the sample, we repeated the analysis without the biased participants. Participants were considered to have a strong response bias, when more than 90% or less than 10% of central sounds were indicated as coming from the right side in auditory-only trials. Based on this criterion, we excluded five subjects, 3 of which had a strong left-ward bias and 2 had a strong right-ward bias (see supplementary material S1 as well as the online [OSF repository](https://osf.io/sa9dn/?view_only=143258c0e5ef4c3f956c3d661c7bd209)).

### Diffusion Modelling

We applied diffusion modeling using fast-dm (Voss & Voss, 2007) to investigate the cognitive mechanisms underlying the odorant-induced localization bias. Accordingly, the analysis focused exclusively on trials in which the auditory stimulus was presented from the central location. Prior to model estimation, RTs faster than 150 ms or outside of the response interval (> 1000 ms) were discarded. Further, to identify outliers, the data was log-transformed and z-standardized within each participant and trials with RTs more than 3 standard deviations from the mean were excluded.

Model parameters were estimated separately for each participant using the maximum likelihood approach. Model specifications were implemented via custom control files and executed in MATLAB using system-level calls to the fast-dm binary. Left button presses were assigned to the lower decision boundary, and right button presses to the upper boundary. Participants with fewer than 4% of trials at one of the decision boundaries were excluded from this analysis(Lerche et al., 2017), leaving 22 subjects for this analysis.

Our primary model allowed both the drift rate (v) and starting point (zr) to vary across the four stimulation conditions (left-nostril, right-nostril, birhinal, and a-only). This combined model was theoretically motivated to directly differentiate whether the observed behavioral bias may arise from either perceptual-level modulation (reflected in drift rate), response-level bias (reflected in starting point), or a combination of both. Boundary separation (a), non-decision time (t₀) and variability in non-decision time (st₀) were freely estimated but fixed across conditions. Variability parameters for drift rate (sv) and starting point (szr) as well as other components (e.g., contaminants, execution time differences) were fixed at zero to ensure stable parameter estimation.

#### Model Fit

To evaluate how well the model accounted for the observed data, we simulated 500 response time distributions per subject and condition using the *construct-samples* routine in *fast-dm*, based on each participant’s estimated parameters. We compared simulated and observed data for each RT quartile (.25, .50, .75) and the proportion of right responses. Model fit was assessed qualitatively through visual inspection of these plots and further supported by computing Pearson correlations between observed and predicted values within each condition across participants. The closer the data points are to the diagonal and the higher the correlations, the better the model fit.

#### Statistical analysis of parameter estimates

Drift rate and starting point were statistically compared between conditions using one-sided paired sample *t*-tests.

Specifically, for each parameter, we conducted separate comparisons between the experimental conditions (right-nostril and left-nostril stimulation) and each of the two control conditions (birhinal and a-only). Building on prior results by Liang et al(Liang et al., 2022), if the localization bias is driven by a perceptual shift, we expected that right-nostril stimulation results in more positive drift rates, whereas left-nostril stimulation results in more negative drift rate (i.e., reflecting a shift in evidence accumulation towards the respective response side). Analogously, if the localization bias is driven by a decision bias, starting point estimates should shift towards the upper decision boundary for right-nostril stimulation and towards the lower decision boundary for left-nostril stimulation.

#### Parameter–Behavior Correlation Analysis

To evaluate whether diffusion model parameters captured the odor-induced bias in auditory localization, we assessed the relationship between model-derived parameter changes and changes in behavioral responses. For each participant, we computed the difference in starting point (zr) and drift rate (v) between each odor condition (right-nostril and left-nostril stimulation) and the two control conditions (birhinal and a-only). These differences were then correlated with the corresponding change in the proportion of rightward responses for the same condition contrasts. Separate Pearson correlation analyses were conducted for left-nostril stimulation vs. control and right-nostril-stimulation vs. control comparisons, using both a-only and birhinal stimulation as a baseline.

This analysis was performed for the model in which both drift rate and starting point varied across conditions (the combined model), as well as for the simpler models in which only one parameter varied (see the *Alternative models* section below). Scatterplots were generated to visualize the relationship between parameter shifts and behavioral bias. This approach allowed us to test whether individual differences in parameter changes (e.g., Δdrift or Δstarting point) systematically predicted the behavioral effect, thereby linking cognitive modeling parameters to observed shifts in the proportion of right-ward responses.

#### Alternative models

While the combined model is the only one that allows us to directly differentiate contributions from drift rate and starting point to the observed odorant-induced sound localization bias, more parsimonious models are possible. Therefore, a supplementary analysis included the estimation of two simpler models: one in which only drift rate varied across conditions, and another in which only starting point varied. All other model parameters were set as described above. To compare model fit of those simpler models with the combined model, we computed the Akaike information criterion (AIC; Akaike, 1973) as well as Akaike weights (Wagenmakers & Farrell, 2004). The statistical comparisons of parameter estimates (drift rate and starting point) between conditions as well as the assessment of the relationship between model-derived parameter changes and changes in behavioral responses were conducted as described for the combined model. Results for these models are briefly summarized in the main text, with full model fit visualizations and parameter estimates presented in the Supplementary Material (supplementary section S4).

### Analysis of alpha power lateralization

To obtain a time-resolved measure of posterior alpha power lateralization, EEG data from four left-hemispheric (i.e., PO7, P7, P3, PO3) and four right-hemispheric (i.e., PO8, P8, P4, PO4) channels were considered (Figure 5A). Alpha power (8-12 Hz) was averaged across electrode sites on the same side as the focus of spatial attention (i.e., ipsilateral) and on the opposite side (i.e. contralateral). To quantify the time-resolved lateralization of alpha power, the alpha lateralization index (ALI) was calculated as follows:

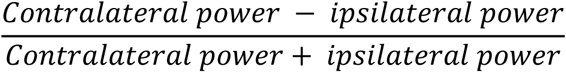

The analysis time-window to derive mean ALI values as input for a rmANOVA was determined based on a collapsed localizer approach(Luck & Gaspelin, 2017). First, the peak in the grand average waveform, collapsed across all conditions and subjects, was determined. Then, a 300 ms window was centered on the respective peak value (i.e., peak ±150 ms). Subsequentially, mean ALI values were obtained per condition and subject.

#### Central Sounds

For the analysis of central sound-trials, the contralateral and ipsilateral electrode sites were assigned relative to the chosen response side (left vs. right). Accordingly, we assumed that the focus of spatial attention shifted to the chosen response hemifield. To account for partially extremely unbalanced trial numbers across response categories, a weighted average was computed when collapsing across trials from different response categories.

First, to verify the presence of a significant lateralization of alpha power in response to a centrally presented sound that is classified as either right- or left-lateralized, irrespective of the odorant-conditions, the grand-average data was tested against zero. To this end, a non-parametric cluster-corrected sign-permutation test was conducted, using the *cluster_test()* and *cluster_test_helper()* functions provided by Wolff and colleagues(Wolff et al., 2017). The *cluster_test_helper()* function generated a null distribution by randomly flipping the sign of the input data of each participant with a probability of 50%. This procedure was repeated 10,000 times. The resulting distribution was submitted to the *cluster_test()* function, which identified those clusters in the actual data that were greater than what would we expected under the null hypothesis. The cluster-forming threshold as well as the cluster significance threshold were set to *p* < .05. Only post-stimulus time-points were considered as input data, since the pre-stimulus period in the full epochs (i.e., -1082 to 1082 ms relative to sound onset) strongly overlapped with the previous trial. Given that the ALI was expected to be negative in case of a shift of attention towards the chosen response hemifield, the cluster-corrected sign-permutation test was one-sided.

Second, to test differences in lateralization magnitude between conditions, mean ALI values were derived and submitted to a one-way rmANOVA including the within-subject factor *stimulation* (congruent, incongruent, birhinal, a-only). The collapsed localizer approach, described above, yielded an analysis time window in-between 389 ms to 689 ms relative to sound onset. Analogous to the behavioral data analysis, congruent odorant-sound stimulation refers to trials in which the chosen response (left vs. right) was in accordance with the stimulated nostril side. Accordingly, incongruent odorant-sound stimulation includes all trials in which the chosen response was opposite to the stimulated nostril side (e.g., left-nostril stimulation and a right-ward response).

#### Lateral Sounds

For the analysis of lateral sounds, only correct responses were included in the analysis. Contralateral and ipsilateral electrode sites were assigned according to the physical sound position. The collapsed localizer approach yielded an analysis time window ranging from 309 to 609 ms post-sound onset. Mean ALI values were submitted to a rmANOVA including the factors *stimulation* (congruent, incongruent, birhinal, a-only) and *sound position* (±15° azimuth, ± 5° azimuth). To follow-up on significant interaction effects, one-sided paired sample *t*-tests were conducted. We expected alpha power lateralization to be increased by a congruent, lateral odorant stimulation or diminished by an incongruent odorant in comparison to controls (birhinal stimulation, a-only). Due to a limited number of trials (i.e., ∼30 trials per cell), the factor *sequence half* (first, second) was omitted from the primary EEG analysis. However, for comparability with the behavioral analysis, a supplementary 3-way rmANOVA was run to check for potential interactions with *sequence half* (see supplementary section, S5).

### Transparency, Openness, and Constraints of Generality

We report how we determined our sample size, all data exclusions, all manipulations, and all measures in the study. All data and code used to generate the present findings will be made publicly available on the [Open Science Framework (OSF)](https://osf.io/sa9dn/?view_only=143258c0e5ef4c3f956c3d661c7bd209) upon acceptance for publication of this manuscript. A view-only link for review is currently available [here](https://osf.io/sa9dn/overview?view_only=143258c0e5ef4c3f956c3d661c7bd209). This study’s design, hypotheses and analyses were not preregistered. Analyses were conducted using JASP (v.0.18.1.0; JASP Team, 2024) as well as custom-written MATLAB (R2023b and R2021b) scripts relying on the open-source toolboxes EEGLAB (v2021.1) and fast-dm (Voss & Voss, 2007). Reflecting the above-described sample characteristics, inclusion criteria, and design parameters, this work is intended to generalize to young healthy adults with intact auditory, olfactory, and trigeminal function. Accordingly, caution is warranted when generalizing the reported findings to older adults, individuals with sensory or neurological impairments, or to more naturalistic listening environments.

## Results

Figure 2 illustrates the proportion of right-ward responses for centrally presented sounds, the proportion of correct responses for lateralized sounds, as well as response times. For easier readability, selected non-significant paired comparison statistics are summarized (e.g., all *p* ≥ 0.282). For full transparency, complete statistical reports of all results are provided in the supplementary material or in the linked [OSF](https://osf.io/sa9dn/?view_only=143258c0e5ef4c3f956c3d661c7bd209) repository.

**Figure 2.**
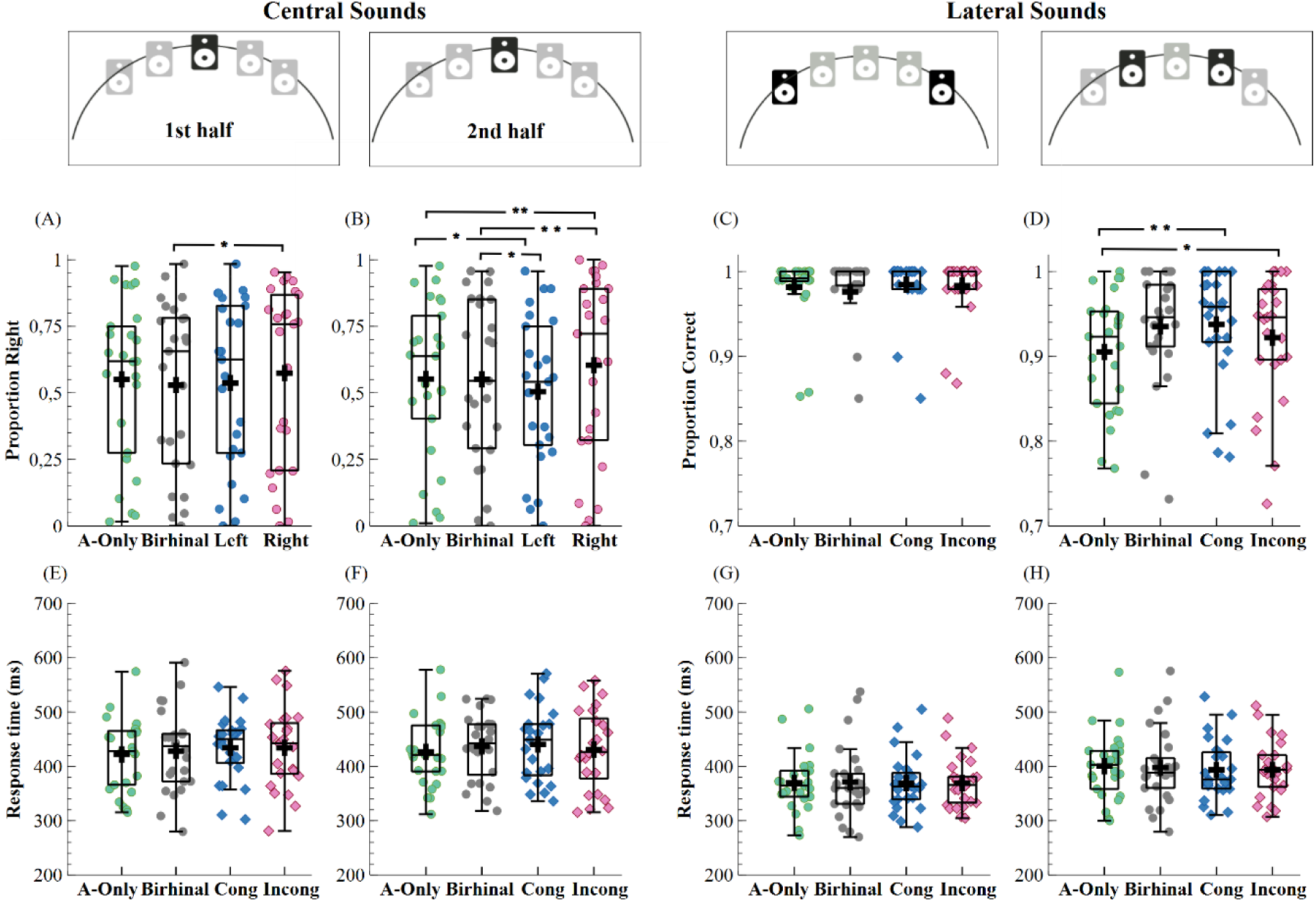
Behavioral Results. The middle panels (A-D) shows the proportion of right-ward responses for centrally presented sounds, separately for the first (A) and second half (B) of the sound sequence, as well as the proportion of correct responses for the lateralized sounds (C-D). The bottom panel (E-H) shows response times for centrally presented sounds (E-F) as well as lateralized sounds (G-H). Boxplots illustrate the distribution of the data, with horizontal lines marking the median, whereas the plus indicates the mean, and whiskers denote +-1.5 IQR. Dots and diamonds show individual participant means. * p < .05, ** p <.01 in one-sided post hoc planned contrasts. For central sounds, congruent (cong) refers to trials in which the stimulated nostril and the chosen response side were compatible (i.e., left nostril stimulation and left-ward response; right-nostril stimulation and right-ward response), whereas incongruent (incong) refers to trials in which they were incompatible (i.e., left-nostril stimulation and right-ward responses; right-nostril stimulation and left-ward response). For lateralized sounds, congruency refers to the side of unilateral odorant stimulation relative to the actual sound position. A-only = Auditory-only.

### Behavioral results

#### After-effect of monorhinal trigeminal stimulation biases sound localization when sound cues are highly ambiguous

A two-way rmANOVA (stimulation × sequence half) on the proportion of right-ward judgements in response to central sounds was conducted to assess the influence of odorant-stimulation on sound localization under conditions of high spatial uncertainty (Figure 2). While there was no significant main effect of sequence halves, F(1,72) = 0.112, p = .741, η_p_^2^ = 0.005, we observed a significant main effect of stimulation, F(3,72) = 4.590, p = .005. η ^2^= 0.161, as well as a significant interaction, F(3,72) = 3.563, p = .018, η_p_ ^2^= 0.129.

To resolve the interaction, we contrasted left and right-nostril stimulation against the control conditions (auditory-only, birhinal stimulation), respectively. Comparisons were conducted separately for trials within the first and second half of the sound sequence.

In the first half of the sequence (Figure 2A), congruent right-nostril stimulation increased the proportion of right-ward judgements compared to the birhinal control condition, t(111.475) = 2.074, p = .020, d = 0.517 [95% CI 0.161 ∞]. The other comparisons in the first half of the sequence were not significant (all p ≥ 0.282, see supplementary table S1). In the second half of the sequence (Figure 2B), left nostril stimulation significantly decreased the proportion of right-ward judgements compared to the birhinal, t(111.475) = -2.153, p = .017, d_z_ = -0.430 [95% CI -∞ -0.081], and auditory-only control conditions, t(111.475) = - 2.199, p = .015, d_z_ = -0.411 [95% CI -∞ -0.064]. In contrast, right-nostril stimulation significantly increased the right-ward indications compared to birhinal stimulation, t(111.475) = 2.441, p = .008, d_z_ = 0.676 [95% CI 0.304 ∞], and compared to the auditory-only control condition, t(111.475) = 2.395, p = .009, d_z_ = 0.378 [95% CI 0.033 ∞]. These findings replicate the odorant-induced sound localization bias, first reported by Liang and colleagues(Liang et al., 2022) and suggests an after-effect rather than an immediate effect of odorant stimulation.

#### Response times for central sounds are not affected by odorant stimulation

An two-way rmANOVA for reaction times in response to the centrally presented sounds did neither render a significant main effect of stimulation, F(3,60) = 0.861, p = .466, η_p_^2^ = 0.41, or of sequence half, F(1,20) = 0.981, p = .334, η_p_ ^2^ = 0.047, nor a significant interaction effect, F(3,60) = 0.399, p = .754 η ^2^ = 0.020. This suggests that response times are not affected by odorant stimulation.

#### Odorant-induced sound localization bias diminishes with increasing spatial discernability of sound cues

To assess whether the observed odorant-induced sound localization bias persists when sound cues become increasingly discernable, we performed a three-way rmANOVA (stimulation × sound position × sequence half) on the proportion of correction responses for lateralized sounds. A significant main effect of sound position, F(1,24) = 34.749, p < .001, η_p_ ^2^= 0.591, confirms the descriptive pattern of better performance for sounds at outer (i.e., +/-15°, M = 98.14%, SD = 4.32%) as opposed to intermediate positions (i.e., +/- 5°, M = 92.48%, SD = 6.54%, see Figure 1-E). Further, a significant main effect of stimulation, F(2.005,48.130) = 3.243, p = 0.048, η_p_ ^2^= 0.119, was obtained after Greenhouse-Geisser correction (approx. χ^2^(5) = 16.613, p = .005, GGε = 0.668). The latter was modulated by sound position, as indicated by a significant interaction between stimulation and sound position, F(3,72) = 5.252, p = .002, η_p_ ^2^= 0.180. None of the effects involving the factor sequence half was significant (all p ≥ .243).

To resolve the significant interaction between stimulation and sound position, planned contrasts were applied to test the effect of stimulation on the localization of sounds at outer (±15° azimuth) and intermediate (± 5° azimuth) speaker positions, respectively. There were no significant differences between the stimulation conditions at outer positions (all p ≥ 0.255, Figure 2C, see supplementary table S2). However, stimulation influenced localization at intermediate positions significantly (Figure 2-D). Specifically, compared to an auditory-only control condition, localization accuracy was significantly higher when chemosensory stimulation and sound position were congruent (i.e., odorant and sound lateralized to same side), t(140.015) = 4.266, p < .001, d_z_ = 0.791 [95% CI 0.406 ∞]. Surprisingly, incongruent stimulation (i.e., odorant presented on the opposite side of the sound source) also significantly increased localization accuracy compared to an auditory-only control condition, t(140.015) = -2.221, p = .028, d_z_ = 0.359 [95% CI -0.050 0.760]. This suggests a generally alerting effect of odorant stimulation; although, compared to birhinal stimulation, performance with incongruent stimulation was slightly decreased, t(140.015) = 1.713, p = .045, d_z_ = 0.306 [95% CI -0.034 ∞].

To support the interpretation of a general altering effect of odorant stimulation, we additionally compared the accuracy of the two control conditions at mid-speaker positions. This data-driven analysis confirmed that birhinal stimulation (M = 93.50%, SD = 6.70%) also improved localization performance compared to auditory-only controls (M = 90.51%, SD = 6.93%), t(140.015) = 3.934, p < .001, d_z_ = 0.566 [95% CI 0.138 0.984].

#### Response times for clearly lateralized sounds are not affected by odorant stimulation

In contrast to the proportion of correct responses, reaction times for lateralized sounds were only marginally affected by the manipulated factors. Mainly, participants responded faster to sounds at outer (+/- 15° azimuth: M = 369.787 ms, SD = 52.16 ms) than intermediate (+/- 5° azimuth: M = 396.423, SD = 55.50 ms) sound positions. Apart from this main effect of sound position, F(1,24) = 142.282, p < .001, η_p_ ^2^=0.856, no further effects reached significance (all p ≥ .084, see supplementary table S3). Note that all results remained unchanged when strongly biased participants (i.e., more than 90% or less than 10% of central sounds in the auditory-only condition were indicated as coming from the right side; n = 5) were excluded from the analysis (see supplementary material section S2).

### Diffusion Modeling

To investigate whether the odor-induced bias in auditory localization reflects a perceptual or a decisional mechanism, we applied a diffusion modeling framework to the data from centrally presented sound trials. The primary analysis includes a *combined model* in which both drift rate (reflecting the rate of evidence accumulation) and starting point (reflecting response bias) were allowed to vary across the four odor conditions (left-nostril, right-nostril, birhinal, a-only). Model parameters were estimated individually for each participant, and model fit was evaluated by comparing predicted and observed RT quantiles and choice proportions.

Across participants and conditions, predicted values closely match the empirical data, as reflected in both graphical assessments (Figure 3) as well as Person correlations (all Pearson r > 0.93), indicating good model fit.

**Figure 3.**
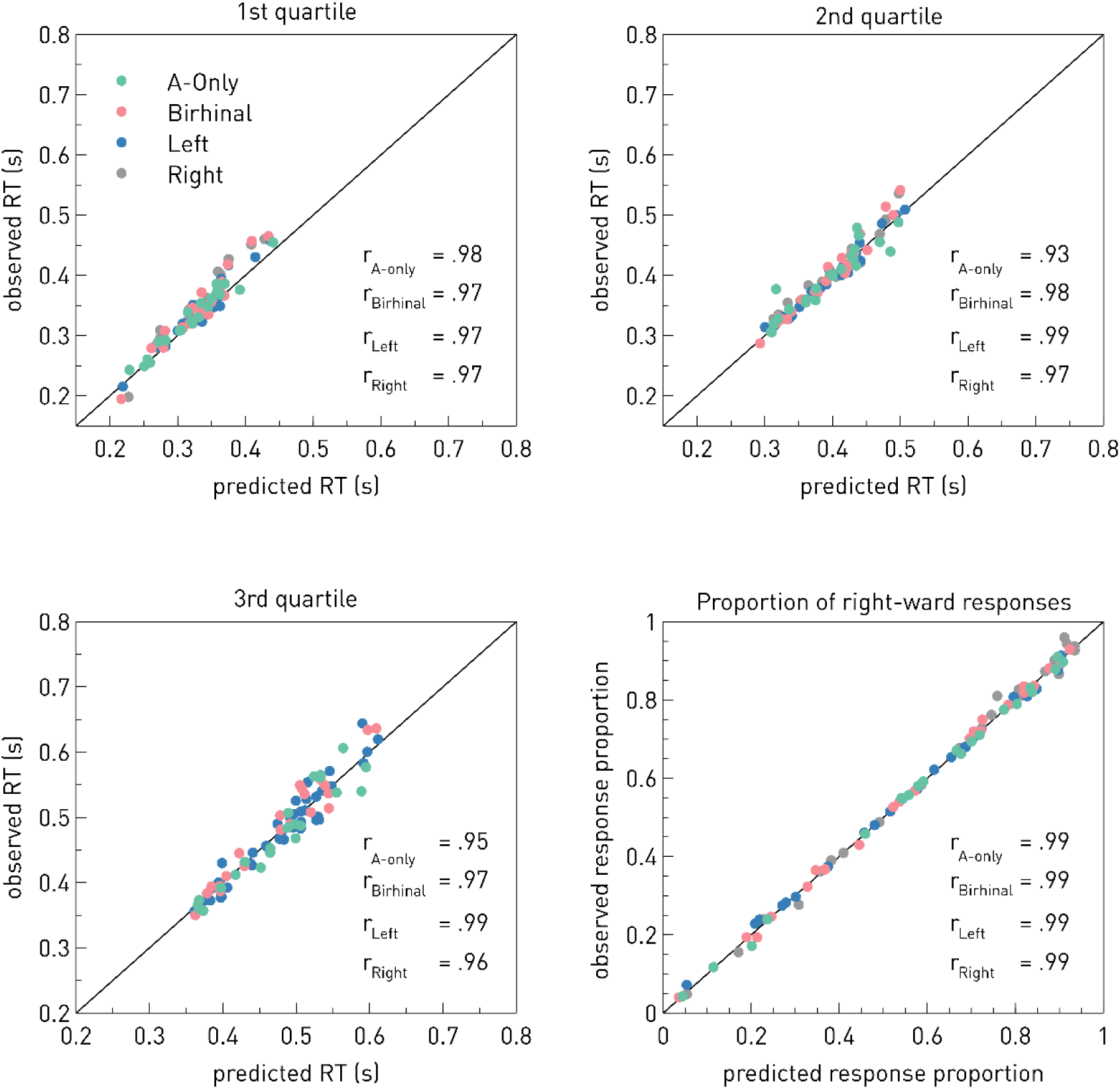
Graphical Model Fit. Scatter plots show the first three quartiles (.25, .5, .75) of the observed response time distribution as well as the observed proportion right-ward responses as a function of the corresponding value from the predicted distribution. Circular data points represent single subject data, separately for each condition. r denotes the corresponding Pearson correlation coefficients.

As shown in Figure 4A, drift rate estimates closely mirrored the behavioral pattern observed in the proportion of rightward responses: drift rate was higher with right-nostril stimulation (*M* = 0.945, *SD* = 1.90) compared to both control conditions (A-only: *M* = 0.404, *SD* = 1.26; Birhinal Odorant: *M* = 0.496, *SD* = 1.56), and lower with left-nostril stimulation (*M* = 0.279, *SD* = 1.49). A one-sided paired-sample *t*-test revealed a non-significant trend towards an increase in drift rate for right-nostril stimulation compared to auditory-only controls, *t*(21) = 2.11, *p* = .024, p_corr_ = .096, *d_z_* = 0.45, 95% CI [0.076, ∞], but not compared to birhinal odorant stimulation, *t*(21) = 1.15, *p* = .131, p_corr_ = .193, *d* = 0.25, 95% CI [–0.113, ∞]. Similarly, drift rate for left-nostril stimulation did not significantly differ from controls (all *p*_corr_ > .193, see supplementary table S6).

**Figure 4.**
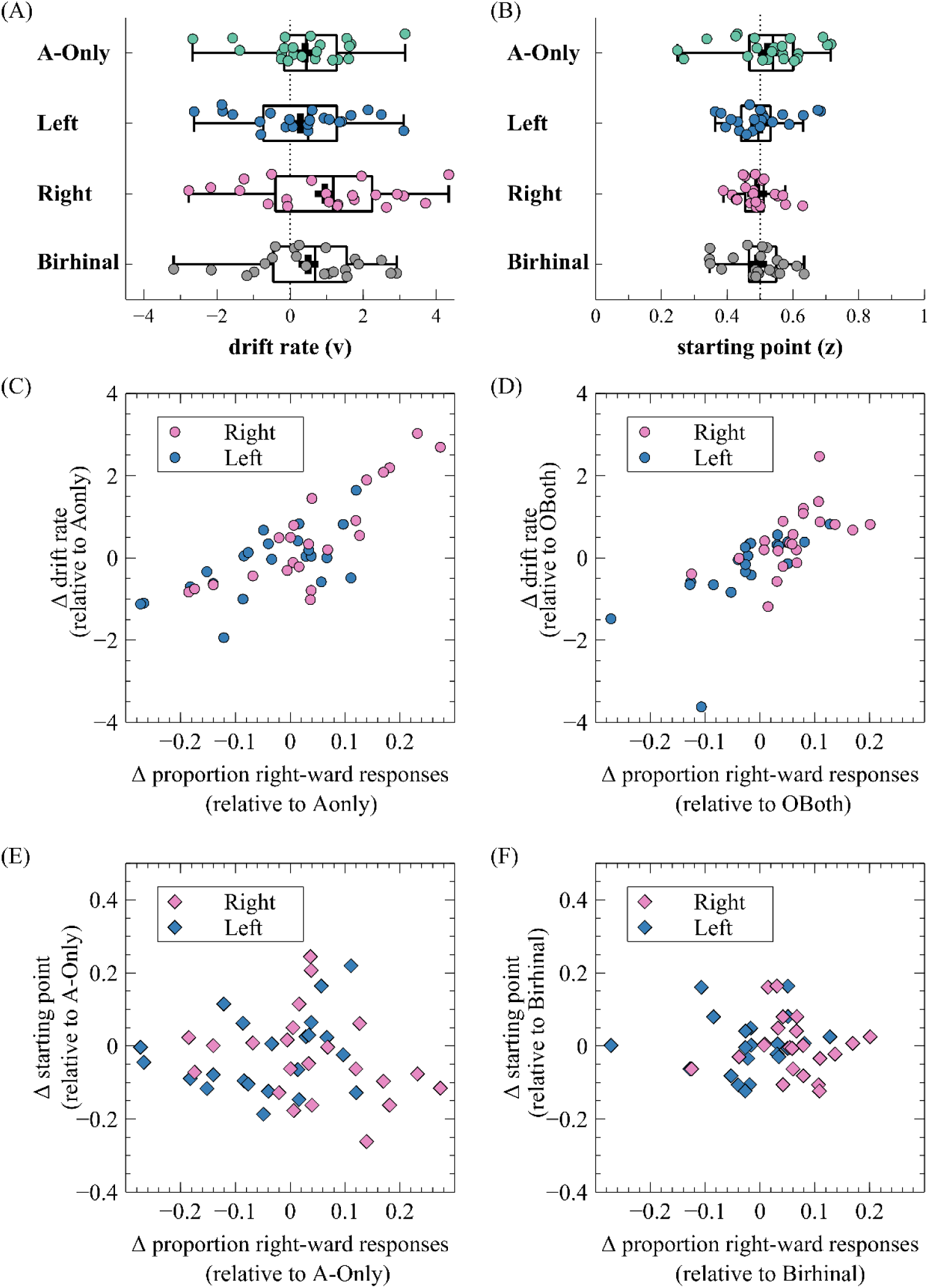
Diffusion Model Estimates for the combined model, allowing both starting point and drift rate to vary between conditions. Panel (A) and (B) depicts boxplots showing the distribution of drift rate and starting point estimates across conditions, respectively. The central line represents the median, the box spans the interquartile range (IQR), and whiskers extend to 1.5 times the IQR. Panels (C) and (D) illustrate the relationship between the relative differences in drift rate and response proportions for Odorant-Left and Odorant-Right trials relative to controls. Analogously, panels (E) and (F) illustrate the relationship between the relative differences in starting point and response proportions for Odorant-Left and Odorant-Right trials relative to controls.

**Figure 5.**
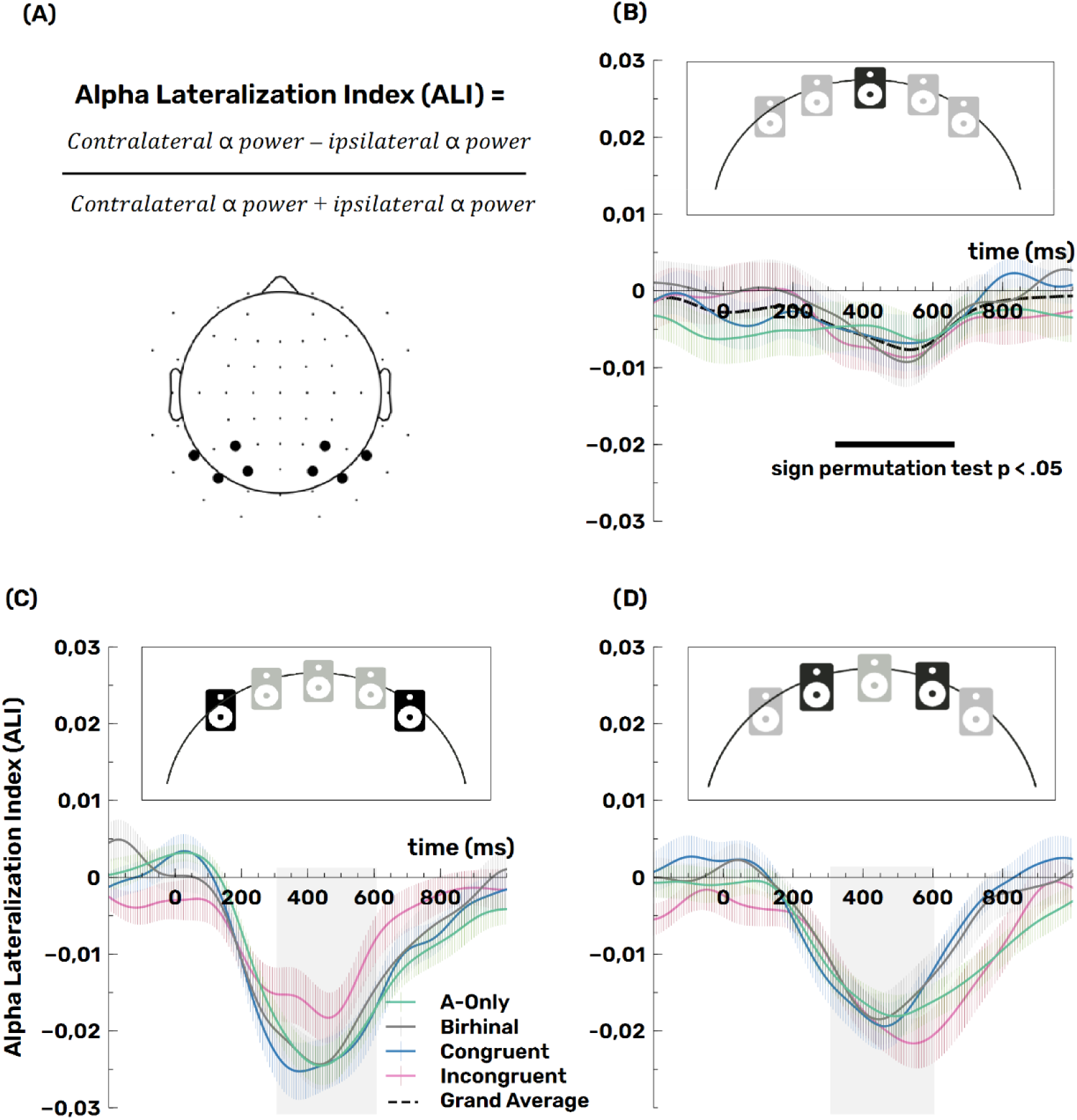
Alpha power lateralization. Magnitude of alpha lateralization is modulated by congruency between chemosensory and auditory spatial information, but only when disparity between the senses is high. (A) Alpha power (8-12 Hz) lateralization was expressed in terms of a normalized hemispheric difference between contralateral and ipsilateral electrode sites. Panels (B)-(D) depict the time course of alpha power lateralization per condition for sounds presented at (B) central, (C) outer (± 15° azimuth), and (D) intermediate (± 5° azimuth) positions, respectively. In the former case, contralateral and ipsilateral alpha power was computed relative to the chosen response category (right vs. left) instead of the physical sound position.

As shown in Figure 4B, starting point estimates showed no systematic modulation across odorant conditions. No statistically significant differences in starting point were found between left-nostril stimulation (*M* = 0.50, *SD* = 0.09) and controls (Birhinal: *M* = 0.49, *SD* = 0.08; A-Only: *M* = 0.52, *SD* = 0.13; all *p* > 0.32) or right-nostril stimulation (*M* = 0.49, SD = 0.06) and controls (all *p_corr_* > 0.64, see supplementary table S7).

Critically, relative differences in drift rate – but not starting point – were significantly associated with odorant-induced shifts in rightward response proportions relative to controls. As shown in Figure 4C, the difference in drift rate between left-nostril stimulation and auditory-only trials was significantly correlated with the corresponding decrease in proportion right responses, *r* = 0.65, *p* = 0.011, *R^2^* = 0.42. Similarly, the drift rate differences between right-nostril stimulation and auditory-only trials were strongly correlated with the respective increase in rightward responses, *r* = 0.84, *p* < .001, *R^2^* = 0.70. A comparable pattern was observed when using birhinal stimulation as the control condition (see Figure 4D, right-nostril: *r* = 0.56, *p* = 0.007, *R^2^* = 0.32; left-nostril: *r* = 0.67, *p* < .001, *R^2^* = 0.46). By contrast, correlations between starting point differences and changes in rightward response proportions were small and non-significant across all comparisons (Figure 4 E-F, all |r|≤ .29, all *p* > .19).

Together, these findings suggest that individual differences in the odorant-induced behavioral bias were more closely associated with changes in drift rate than with shifts in starting point, suggesting a modulation of perceptual evidence rather than decisional bias. To evaluate whether this interpretation holds when accounting for model parsimony, we next compared the combined model to two simpler model structures: one in which only drift rate varied across conditions, and one in which only starting point varied. Model comparisons based on the Akaike information criterion (AIC(Akaike, 1998)) and Akaike weights(Wagenmakers & Farrell, 2004) were used to assess relative model fit across participants.

Table 2 provides the mean AIC values for each specified model and the proportion of subjects whose data was best described by the model. The starting point–only model provided the best fit for the largest number of participants (*n* = 15), followed by the drift rate–only model (*n* = 7). The combined model did not outperform the simpler models for any participant. Mean AIC values further reflected this pattern: the starting point model had the lowest average AIC (*M* = 419.76), followed by the drift rate model (*M* = 426.47), and the combined model showed the highest mean AIC (*M* = 437.80). These results suggest that, from a model parsimony standpoint, the simpler models were preferred over the combined model.

**Table 2.** Mean Akaike information criterion (AIC) for each specified model and percentage of the sample whose data were best described by the model. v = drift rate; z = starting point.

| Model | Model parameter varying across conditions | Number of estimated parameters | Mean AIC | Best fitting model for % of participants |
| --- | --- | --- | --- | --- |
| Star-Drift-Model | v, z | 11 | 437.80 | 0% |
| Drift-Model | v | 8 | 426.47 | 32% |
| Starting Point Model | z | 8 | 419.76 | 68% |

Interestingly, the split in best-fitting models – drift-only for some participants and starting point–only for others – may reflect individual differences in the cognitive mechanisms underlying the odorant-induced bias, with some individuals showing more perceptual (drift) and others more decisional (starting point) contributions.

### EEG Results: Alpha power lateralization

#### Alpha power lateralization in response to central sounds confirms that participants allocate attention to the chosen sound location, but remains insensitive to condition differences

Figure 2 (B-D) depicts the alpha lateralization index (ALI) for each stimulation condition (auditory-only, birhinal, congruent, incongruent) as a function of sound position (central vs. outer vs. intermediate). For centrally presented sounds, the alpha lateralization index is computed relative to the given response such that congruency between odorant-stimulation and the perceived (or chosen) sound location is reflected. A cluster-corrected sign-permutation test showed a small, but significant (p = .0395) overall lateralization of alpha power in-between ∼320 to 650 ms post-stimulus onset, indicative of a spatial shift of attention towards the selected hemifield. However, a rmANOVA of average alpha lateralization magnitude (revealed no significant differences between the stimulation conditions, F(2.06, 49.34) = 0.225, p = .805, η_p_ ^2^= 0.009, after Greenhouse-Geisser correction (ε = 0.69).

#### Alpha power lateralization in response to lateral sounds indicates a disruption of spatial attention when disparity between auditory and chemosensory cues is strong

A two-way rmANOVA revealed no significant main effect of *stimulation*, *F*(3,72) = 0.799, *p* = .498, *η*_p_ ^2^= 0.032. In contrast, a significant main effect of *sound position* was evident, *F*(1,24) = 6.976, *p* = .014, *η*_p_ ^2^= 0.225, in addition to a significant interaction of *stimulation* and *sound position*, *F*(3,72) = 3.434, *p* = .021, *η*_p_ ^2^= 0.125. Follow-up one-sided paired-sample *t*-tests showed that no significant differences between conditions were obtained at lateralized positions close to midline (± 5° azimuth); all *p_corr_* ≥ .64, see supplementary table S8), while alpha power lateralization was significantly modulated by odorant-stimulation at outer sound positions (±15° azimuth). Specifically, incongruent odorant stimulation (*M* = -0.015, *SD* = 0.0149) resulted in a reduction of alpha power lateralization compared the a-only condition (*M* = -0.022, *SD* = 0.011), *t*(24) = 2.9165, *p* = .0038, *p_corr_* = 0.0189, *d_z_* = 0.583, 95% CI [0.221 ∞], as well as birhinal stimulation (*M* = -0.021, *SD* = 0.016), *t*(24) = 2.4122, *p* = .012, *p_corr_* = .0398, *d_z_* = 0.482, 95% CI [0.130 ∞]. The magnitude of alpha power lateralization at outer sound positions in congruent trials did not differ significantly from the control conditions with birhinal stimulation *t*(24) = -0.8104, *p* = .426, *p*_corr_ = .715, *d_z_* = -0.162, 95% CI [-∞ 0.171], or a-only stimulation, *t*(24) = -0.5731, *p* = .5719, *p*_corr_ = .715, *d_z_* = -0.115, 95% CI [- ∞ 0.217]. Lastly, incongruent odorant stimulation (*M* = -0.015, *SD* = 0.0149) resulted in a reduction of alpha power lateralization compared to congruent stimulation (*M* = -0.023, *SD* = 0.0169), *t*(24) = -3.0827, *p* = .0025, *p_corr_* = 0.019, *d_z_* = -0.617, 95% CI [-∞ -0.251]. A supplementary analysis, including the additional factor *sequence half,* confirmed the two-way interaction between sound position and condition, *F*(3,72) = 2.904, *p* = .041, *η*_p_^2^= 0.108, while the three-way interaction of *sound position, condition, and sequence half* was not significant, *F*(3, 72) = 1.323, *p* = .274, *η*_p_ ^2^= 0.052 (see supplementary table S9).

## Discussion

Reconciling conflicting input from different sensory channels poses a unique challenge for the brain. Here, we investigated to what extent a task-irrelevant, yet highly salient (i.e., trigeminally potent) bimodal odorant affects two-alternative forced-choice sound localization judgements. We show that the proportion of centrally presented sounds that was categorized as coming from the right increased with right-nostril odorant stimulation, while it decreased with left-nostril stimulation (compared to two control conditions with birhinal nostril and auditory-only stimulation, respectively). The results replicate previous findings of an odorant-induced sound localization bias (Liang et al., 2022).

Notably, the odorant-induced localization bias was only consistently present in trials during the second half of the sound sequence, speaking in favor of an “after-effect” as opposed to an immediate effect of odorant stimulation. This is in line with the finding that chemosensory processing is relatively slow compared to other sensory modalities (Pause et al., 1997). In particular, trigeminal processing unfolds slowly and can outlast the physical stimulus by several seconds in both neural (Carlson et al., 2013) and perceptual responses (Hummel et al., 1992). Thus, since previous studies have shown that pure olfactory cues cannot be consciously localized, it is most likely that the present effects are driven by the trigeminal component (but see Moessnang et al., 2011 for evidence in favor of residual ability of directional smelling based on purely olfactory cues).

In contrast to the effect observed for centrally presented sounds, the effect of odorant stimulation became spatially unspecific with increasing discernability of the sound cues (i.e., at +/- 5° azimuth). That is, the participants showed better performance in all odorant conditions compared to an auditory-only control condition, irrespective of whether the nostril-side was congruent or incongruent with the actual sound location. This suggests a more general alerting effect (La Buissonnière-Ariza et al., 2012; Michael et al., 2005), in line with the proposed warning function of the trigeminal system (Bessac & Jordt, 2008; Brüning et al., 2014; Terrier et al., 2022). Notably, though, when sounds were most strongly lateralized and hence, clearly discernable in terms of their spatial position (at +/- 15° azimuth), performance was close to ceiling in all conditions and there was no effect of odorant stimulation. Overall, this aligns with the notion that chemosensory cues exert an effect on perception in other modalities only when the latter is ambiguous. Accordingly, Zhou and Chen (2009) found that fearful sweat biased female participants toward rating ambiguous facial expressions as more fearful, while the same odorants had no effect when the facial expression was more apparent. On a similar note, Zhou and colleagues (2010) reported an effect of olfactory cues on binocular rivalry, a form of multistable perception caused by the presentation of dissimilar images to the two eyes (Blake & O’Shea, 2017).

More generally, these results align with the principle of inverse effectiveness in multisensory integration, which posits that the benefit of combining sensory inputs across modalities is strongest when individual inputs are weak or ambiguous (Meredith & Stein, 1986; Stein & Stanford, 2008). Accordingly, Liang et al (2022) argued that the absence of a sound localization bias towards the odorant at sound azimuths greater than 10° supports their interpretation of odorant-sound integration. However, while the behavioral pattern is consistent with this principle, it does not in itself confirm perceptually driven cross-modal integration. Alternatively, the effect may reflect a post-perceptual decision bias such that observers are more inclined to respond in accordance with the stimulation nostril side whenever they are uncertain about the sound location, without an actual change in the perceived spatial position. Notably, even in the absence of clear spatial information (i.e., for central sounds in the auditory-only condition), the majority of participants in the present study exhibited a tendency towards one response side (see also Liang et al., 2022 for a similar effect in some participants). This suggests that participants may have adopted a certain strategy, favoring one side when uncertain. Crucially, however, the odorant-induced localization biased emerged on top of this baseline tendency: the proportion of “right” responses shifted systematically with unilateral odor stimulation relative to these control conditions. Thus, the observed effect cannot be solely attributed to a general response-side preference, leaving the critical question of whether the odor cues did influence perception or simply modulated decision thresholds. To address this, we employed a drift diffusion model approach, allowing us to formally distinguish changes in evidence accumulation (perceptual bias) from shifts in starting point (decision bias) (Voss et al., 2008) The primary model of interest allowed both starting point and drift rate to vary between conditions.

While group-level statistical comparisons of drift rate estimates between conditions revealed no robust effects, the overall pattern of drift rate changes descriptively mirrored the behavioral effects in proportion right responses (see Figure 4-A). Corroborating this observation, our correlational analysis provided compelling evidence, linking changes in drift rate to changes in condition differences in the behavioral reports: Participants with a stronger increase in drift rate in Odorant-Right trials relative to controls showed a significantly stronger increase in the proportion of right responses (relative to controls), and vice versa for Odorant-Left trials (see Figure 4-C and D). This pattern held across both control conditions (A-Only and Odor-Both) and was not observed for changes in starting points. Although the paired comparisons offered limited statistical support, the strength and directionality of the correlations, considering relative condition differences, suggest that drift rate captures meaningful variation in perceptual evidence accumulation, modulated by the odorant stimulation.

By contrast, starting point estimates did reflect individual decision tendencies, such that in participants who were generally more biased towards left responses (even in the control conditions), the starting point was shifted towards the lower decision boundary, while in those with a stronger bias towards the right, the starting point was shifted towards the upper decision boundary. However, these estimates showed no systematic modulation by experimental conditions (Figure 4-E and F). This indicates that while variation in starting point may account for general response tendencies, it does not explain the odorant-induced localization bias. Together these findings support the interpretation that bimodal odorant stimulation influenced the accumulation of sensory evidence, rather than simply altering response strategy.

At the same time, a few cautionary notes are warranted. We further compared the full model (in which both drift rate and starting point were allowed to vary across conditions) to reduced models, allowing only one parameter to vary. Based on AIC values and Akaike weights, the starting point model yielded the best fit for a majority of participants (*n* = 15), whereas the drift rate model showed a better fit in a smaller subset of participants (*n* = 7). Given that model fit indices value parsimony, it is perhaps not surprising that the reduced models outperformed the full model. Interestingly, repeating the correlation analysis in the reduced models revealed that the parameter which was allowed to vary between conditions predicted the relative shifts in rightward response proportions (see supplementary figure S2 and S4, panels F and G). This raises the possibility that both perceptual and decision-level mechanisms may contribute to the observed odorant-induced bias. However, it is important to note that in models where only a single parameter is free to vary, that parameter is more likely to absorb all between-condition variance by design. As such, the combined model, despite being more complex, arguably provides a more informative account, linking individual differences in drift rate to behavior in a way that the starting point parameter did not. In sum, these findings suggest that a perceptual influence on evidence accumulation is likely, but strategic decision-level biases cannot be entirely ruled out and may also play a role in some individuals.

At the electrophysiological level, we observed that EEG alpha-band activity lateralizes toward the hemifield where the sound was localized. Asymmetric modulations of alpha power have long been associated with the allocation of spatial attention (Klatt et al., 2018a, 2018b, 2020; Wöstmann et al., 2018, 2019). Thus, the results are consistent with attentional orienting towards the perceived sound location. However, alpha lateralization for central sounds did not differ between congruent and incongruent odorant stimulation, prompting questions about the relationship between alpha oscillations, sensitivity and attentional gain control. The fact that changes in drift rate (i.e., the quality of perceptual evidence accumulation) track the behavioral sound localization bias towards concurrent odorants aligns with the view that spatial attention enhances sensory gain, thereby improving sensitivity and the quality of information extracted at attended locations (Hillyard et al., 1998). Accordingly, we hypothesized that alpha lateralization should be boosted by the presence of congruent odorant stimulation or diminished in case of an incongruent odorant. On the contrary, alpha lateralization in response to central sounds did not differ between trials that were paired with a spatially congruent as opposed to an incongruent odorant. This rather suggests that the attentional modulation of alpha power is more closely linked to down-stream gating of information, in line with recent proposals (Jensen, 2024). However, methodological factors may have also contributed to the lack of a distinction between congruent and incongruent odor stimulation trials in the alpha power modulations. That is, trial averages may include trials in which the chemosensory cues effectively captured attention, but also trials in which they did not. Including a central response option in future studies may help disambiguate true perceptual shifts (i.e. illusion trials) from trials, unaffected by cross-modal interactions (non-illusion trials) (Bonath et al., 2007; Bruns & Röder, 2010). Further, separating trials by sequence half may have helped to isolate alpha lateralization effects specifically in trials in which a true perceptual shift was more likely. However, limited trial numbers constrained the statistical power of such analyses (supplementary material S2).

In contrast, for less ambiguous, strongly lateralized sounds (+/-15° azimuth), a diminished lateralization of alpha power emerged when the chemosensory cues and the sound location were maximally discrepant (i.e., incongruent) compared to congruent stimulation. In such cases, the trigeminal component of the odorant may have most effectively exogenously captured spatial attention – a phenomenon supported by classical cueing studies, showing that task-irrelevant olfactory cues can attract spatial attention (Ischer et al., 2021, Wudarczyk et al., 2016; albeit mixed evidence exists: La Buissonnière-Ariza et al., 2012). This suggests that large cross-modal disparities interfered with attentional selection, although this disruption was not sufficiently strong to be reflected on the behavioral level, where performance was close to ceiling in all condition.

## Conclusion

Taken together, the present study generalizes previous findings of an odorant-induced sound localization bias to a novel, EEG-compatible paradigm using isopropanol as a bimodal stimulus. By combining behavioral analyses, diffusion modeling, and electrophysiology, we provide converging evidence that the effect is perceptual in nature, driven by changes in the quality of evidence accumulation rather than mere shifts in decisional bias. At the neural level, alpha-band activity lateralized toward the reported sound location and was selectively disrupted under high cross-modal conflict, highlighting the susceptibility of spatial attention to interference from chemosensory input. These findings not only extend the spatial ventriloquism phenomenon to the chemosensory domain but also point to important future questions about how attention and integration mechanisms resolve spatial information across disparate reference frames. More broadly, our approach illustrates the value of combining computational modeling with neural and behavioral data to disentangle the mechanisms underlying cross-modal interactions.

## Authors’ contributions (CRediT)

Conceptualization: C.H. (equal) and L.I.K. (equal); Methodology: C.H. (equal) and L.I.K. (equal); Formal Analysis: C.H. (support) and L. I.K. (lead); Investigation: C.H. (equal) and L.I.K. (equal); Data Curation: C.H. (equal) and L.I.K. (equal); Writing – Original Draft: C.H. (support) and L.I.K. (lead), Writing – Review & Editing: C.H. (support) and L.I.K. (lead); Visualization: C.H. (equal) and L.I.K. (equal)

## Acknowledgements

We would like to thank J. Reiser, A. Hijazi, as well as S. Kattan for their help with EEG data collection. In addition, we would like to thank M. Porta and J. Reinders for the preparation of isopropanol samples as well as N. Koschmieder for assistance with spirometry data acquisition. Finally, we are grateful for S. Getzmann’s and C. van Thriel’s input in discussions concerning the task design.

## Supplementary Material

### S1 Supplementary Behavioral Results: Full statistical reports for the analysis of the proportion of rightward responses for central sounds

Non-significant statistical reports that were abbreviated in the main text are provided here (as well as on OSF) for full transparency.

**Table S1.** Planned contrast results for the proportion right data for central sounds.

|  | **T** | **DF** | **P** | **D<sub>z</sub>** | **95% CI** |
| --- | --- | --- | --- | --- | --- |
| Left-nostril vs. Birhinal, first half | 0.349 | 111.475 | 0.728 | 0.110 | [-0.284 0.502] |
| Left-nostril vs. A-only, first half | -0.645 | 111.475 | 0.260<sup>a</sup> | -0.141 | [ -∞ 0.191] |
| Left-nostril vs. Birhinal, second half | -2.153 | 111.475 | 0.017<sup>a</sup> | -0.430 | [ -∞ -0.081] |
| Left-nostril vs. A-only, second half | -2.199 | 111.475 | 0.015<sup>a</sup> | -0.411 | [ -∞ -0.064] |
| Right-nostril vs. Birhinal, first half | 2.074 | 111.475 | 0.020<sup>a</sup> | 0.517 | [0.161 ∞ ] |
| Right-nostril vs. A-only, first half | 1.081 | 111.475 | 0.141<sup>a</sup> | 0.196 | [-0.138 ∞ ] |
| Right-nostril vs. Birhinal, second half | 2.441 | 111.475 | 0.008<sup>a</sup> | 0.676 | [0.304 ∞ ] |
| Right-nostril vs. A-only, second half | 2.395 | 111.475 | 0.009<sup>a</sup> | 0.378 | [0.033 ∞ ] |

Note that planned contrasts were computed using the pooled error term. <sup>a</sup> denotes one-sided *p*-values. That is, as JASP does not offer one-sided contrasts, the *p*-value was divided by two if the difference in means was in line with the direction of our hypothesis. *d<sub>z</sub>* values were computed using the standard deviation of the condition differences as the denominator.

**Table S2.** Planned contrast results for the proportion of correct responses at lateralized sound positions.

|  | **T** | **DF** | **P** | **D<sub>z</sub>** | **95% CI** |
| --- | --- | --- | --- | --- | --- |
| Congruent vs. Birhinal, outer | 1.143 | 140.015 | 0.128<sup>a</sup> | 0.244 | [-0.092 ∞ ] |
| Congruent vs. A-only, outer | 0.395 | 140.015 | 0.347<sup>a</sup> | 0.264 | [-0.059 ∞ ] |
| Incongruent vs. Birhinal, outer | -0.993 | 140.015 | 0.323 | 0.191 | [-0.207 0.584] |
| Incongruent vs. A-only, outer | -0.245 | 140.015 | 0.807 | 0.126 | [-0.269 0.518] |
| Congruent vs. Birhinal, intermediate | 0.332 | 140.015 | 0.371<sup>a</sup> | 0.058 | [-0.272 ∞ ] |
| Congruent vs. A-only, intermediate | 4.266 | 140.015 | <.001<sup>a</sup> | 0.791 | [0.406 ∞ ] |
| Incongruent vs. Birhinal, intermediate | 1.713 | 140.015 | 0.045<sup>a</sup> | 0.306 | [-0.034 ∞ ] |
| Incongruent vs. A-only, intermediate | -2.221 | 140.015 | 0.028 | -0.359 | [-0.050 0.760] |

Note that planned contrasts were computed using the pooled error term. <sup>a</sup> denotes one-sided *p*-values. That is, as JASP does not offer one-sided contrasts, the *p*-value was divided by two if the difference in means was in line with the direction of our hypothesis. Outer and immediate sound positions refer to sounds played at +/- 15° and +/- 5° azimuth, respectively. *d<sub>z</sub>* values were computed using the standard deviation of the condition differences as the denominator.

**Table S3.** 4 × 2 × 2 rmANOVA (odorant condition × sound position × sequence half) response time analysis at lateralized sound positions.

|  | **DF** | **F** | **P** | **N<sub>p</sub><sup>2</sup>** |
| --- | --- | --- | --- | --- |
| Odorant condition | [2.18, 52.32] | 0.203 | 0.835 | 0.008 |
| Sound position | [1, 24] | 142.282 | < .001 | 0.856 |
| Sequence half | [1, 24] | 3.255 | 0.084 | 0.119 |
| Odorant condition x sound position | [3, 72] | 0.715 | 0.546 | 0.029 |
| Odorant condition x sequence halves | [3, 72] | 0.153 | 0.927 | 0.006 |
| Sound position x sequence halves | [1, 24] | 0.116 | 0.737 | 0.005 |
| Odorant condition x sound position x sequence halves | [2.012, 48.29] | 0.310 | 0.818 | 0.013 |

### S2 Supplementary Behavioral Analyses: Excluding strongly biased participants

As in Liang et al., a subset of participants showed strong response tendencies already in the auditory-only control condition. Participants were considered to have a strong response bias, when more than 90% or less than 10% of mid sounds in the auditory-only condition were indicated as coming from the right side. Based on this criterion, we excluded five subjects, 3 of which had a strong left-ward bias and 2 had a strong right-ward bias. Here, we report only the relevant analyses for central sounds and the proportion of right-ward responses, as these were the ones yielding significant results in the main analysis. For all other corresponding analyses, please see the jasp files in the OSF repository.

**Table S4.** 4 × 2 rmANOVA results for the proportion right data at central sound locations, excluding strongly biased participants.

|  | DF | F | P | N<sub>p</sub><sup>2</sup> |
| --- | --- | --- | --- | --- |
| Odorant condition | [3, 57] | 5.010 | .004** | 0.209 |
| Sequence half | [1, 19] | 0.129 | .723 | 0.007 |
| Odorant condition x sequence half | [3, 57] | 4.117 | .010* | 0.178 |

**Table S5.** Planned contrast results for the proportion right data at central sound locations, excluding strongly biased participants.

|  | T | DF | P | D<sub>z</sub> | 95% CI |
| --- | --- | --- | --- | --- | --- |
| Left-nostril vs. Birhinal, first half | 0.304 | 87.528 | 0.762 | 0.107 | [-0.334 0.545] |
| Left-nostril vs. A-only, first half | -0.603 | 87.528 | 0.274<sup>a</sup> | -0.142 | [ -∞ 0.229 ] |
| Left-nostril vs. Birhinal, second half | -2.216 | 87.528 | 0.015<sup>a</sup> | -0.496 | [ -∞ -0.099] |
| Left-nostril vs. A-only, second half | -2.191 | 87.528 | 0.016<sup>a</sup> | -0.457 | [ -∞ -0.064] |
| Right-nostril vs. Birhinal, first half | 2.057 | 87.528 | 0.022<sup>a</sup> | 0.580 | [0.174 ∞ ] |
| Right-nostril vs. A-only, first half | 1.149 | 87.528 | 0.127<sup>a</sup> | 0.110 | [-0.148 ∞ ] |
| Right-nostril vs. Birhinal, second half | 2.662 | 87.528 | 0.005<sup>a</sup> | 0.894 | [0.447 ∞ ] |
| Right-nostril vs. A-only, second half | 2.686 | 87.528 | 0.005<sup>a</sup> | 0.477 | [0.083 ∞ ] |

Note that planned contrasts were computed using the pooled error term. <sup>a</sup> denotes one-sided p-values. That is, as JASP does not offer one-sided contrasts, the *p*-value was divided by two if the difference in means was in line with the direction of our hypothesis. *d<sub>z</sub>* values were computed using the standard deviation of the condition differences as the denominator.

### S3 Supplementary Results: Combined Diffusion Model, allowing both starting point and drift rate to vary between conditions

**Table S6.** Full statistical results for one-sided paired comparisons, contrasting drift rate for right- and left-nostril stimulation with controls.

|  | **T** | **DF** | **P** | **P<sub>CORR</sub>** | **D<sub>Z</sub>** | **95% CI** |
| --- | --- | --- | --- | --- | --- | --- |
| Left-nostril vs. A-only | -0.736 | 21 | 0.235 | 0.235 | -0.157 | [-∞ 0.198] |
| Left-nostril vs. Birhinal | -1.087 | 21 | 0.145 | 0.193 | -0.232 | [-∞ 0.126] |
| Right-nostril vs. A-only | 2.109 | 21 | 0.024 | 0.096 | 0.450 | [0.07 ∞] |
| Right-nostril vs. Birhinal | 1.152 | 21 | 0.131 | 0.193 | 0.246 | [-0.113 ∞] |

**Table S7.** Full statistical results for one-sided paired comparisons, contrasting starting point for right- and left-nostril stimulation with controls.

|  | **T** | **DF** | **P** | **P<sub>CORR</sub>** | **D<sub>Z</sub>** | **95% CI** |
| --- | --- | --- | --- | --- | --- | --- |
| Left-nostril vs. A-only | -1.013 | 21 | 0.161 | 0.644 | -0.216 | [-∞ 0.141] |
| Left-nostril vs. Birhinal | 0.451 | 21 | 0.672 | 0.883 | 0.096 | [-∞ 0.446] |
| Right-nostril vs. A-only | -1.224 | 21 | 0.883 | 0.883 | -0.261 | [-0.615 ∞] |
| Right-nostril vs. Birhinal | -0.068 | 21 | 0.527 | 0.883 | -0.015 | [-0.365 ∞] |

### S4 Alternative Drift Diffusion Models

In addition to the combined model, we estimated a model in which only starting point (or only drift rate) was allowed to vary between conditions. Below, we present the graphical model fit matrices, parameter estimates and the relationship between changes in starting point (or drift rate) and changes in right-ward response proportions relative to controls.

**Figure S1.**
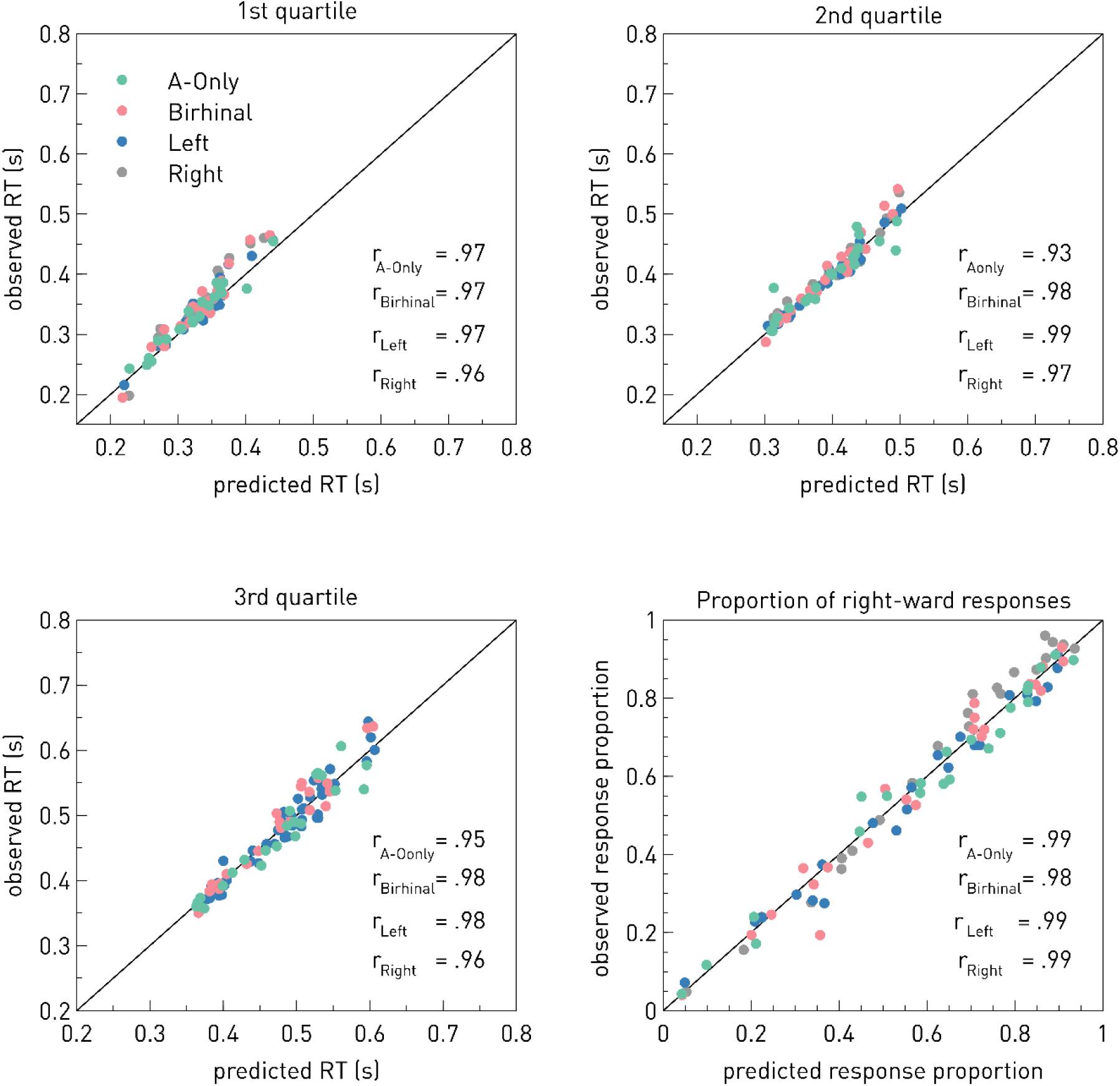
Graphical Model Fit for the “Starting Point Model”, allowing only starting point to vary across odorant conditions. Scatter plots show the first three quartiles (.25, .5, .75) of the observed response time distribution as well as the observed proportion right-ward responses as a function of the corresponding value from the predicted distribution. Circular data points represent single subject data, separately for each condition. r denotes the corresponding Pearson correlation coefficients.

**Figure S2.**
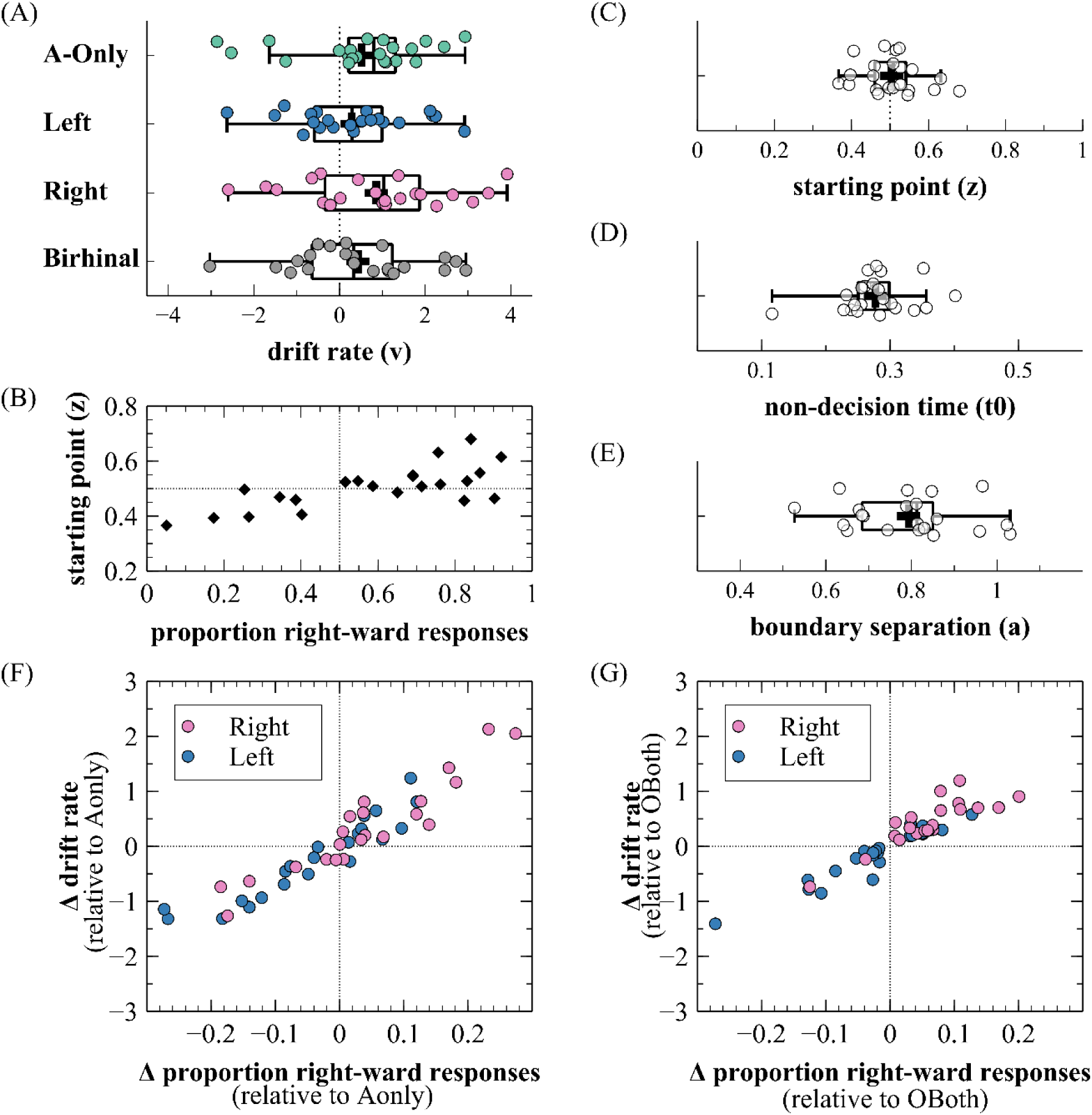
Diffusion Model Estimates for the Starting Point Model, allowing only starting point to vary between conditions. Panel (A) depicts boxplots showing the distribution of starting point estimates across conditions. The central line represents the median, the box spans the interquartile range (IQR), and whiskers extend to 1.5 times the IQR. Panel (B) illustrates the relationship between drift rate estimates and the proportion of right-ward responses. Participants with response proportions below 0.5 tend to have a negative drift rate, whereas participants with response proportions above 0.5 tend to have positive drift rates, illustrating that the model generally captures the pattern of behavioral responses. Panels (C), (D), and (E) depicts boxplots showing the distribution of drift rate(v), non-decision time(t0) and boundary separation (a), all of which were not allowed to vary between conditions. Panels (F) and (G) illustrate the relationship between the relative differences in starting point and response proportions for Odor-Left and Odor-Right trials relative to controls.

**Figure S3.**
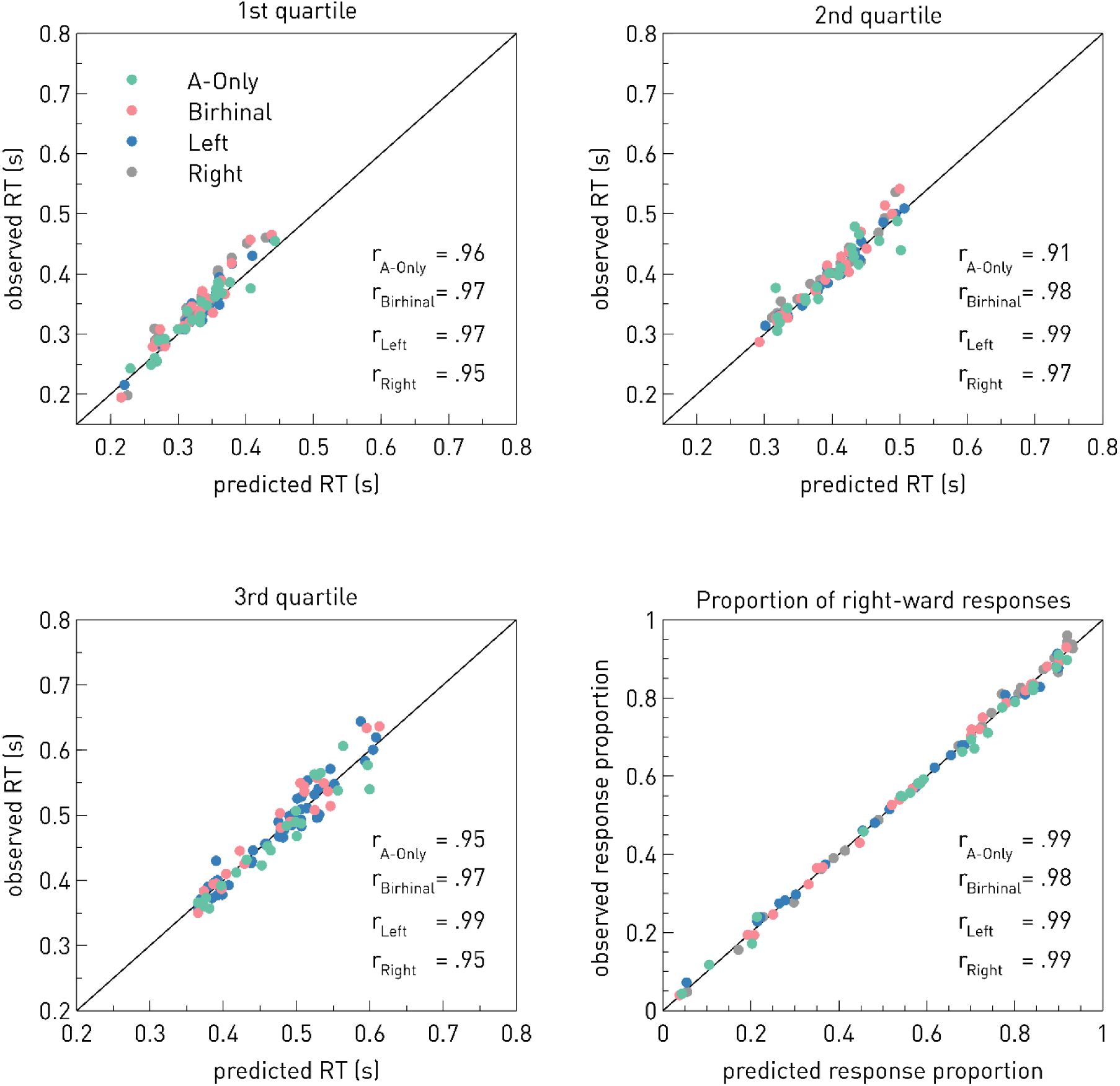
Graphical Model Fit for the “Drift rate model”, allowing only drift rate (v) to vary across odorant conditions. Scatter plots show the first three quartiles (.25, .5, .75) of the observed response time distribution as well as the observed proportion right-ward responses as a function of the corresponding value from the predicted distribution. Circular data points represent single subject data, separately for each condition. r denotes the corresponding Pearson correlation coefficients.

**Figure S4.**
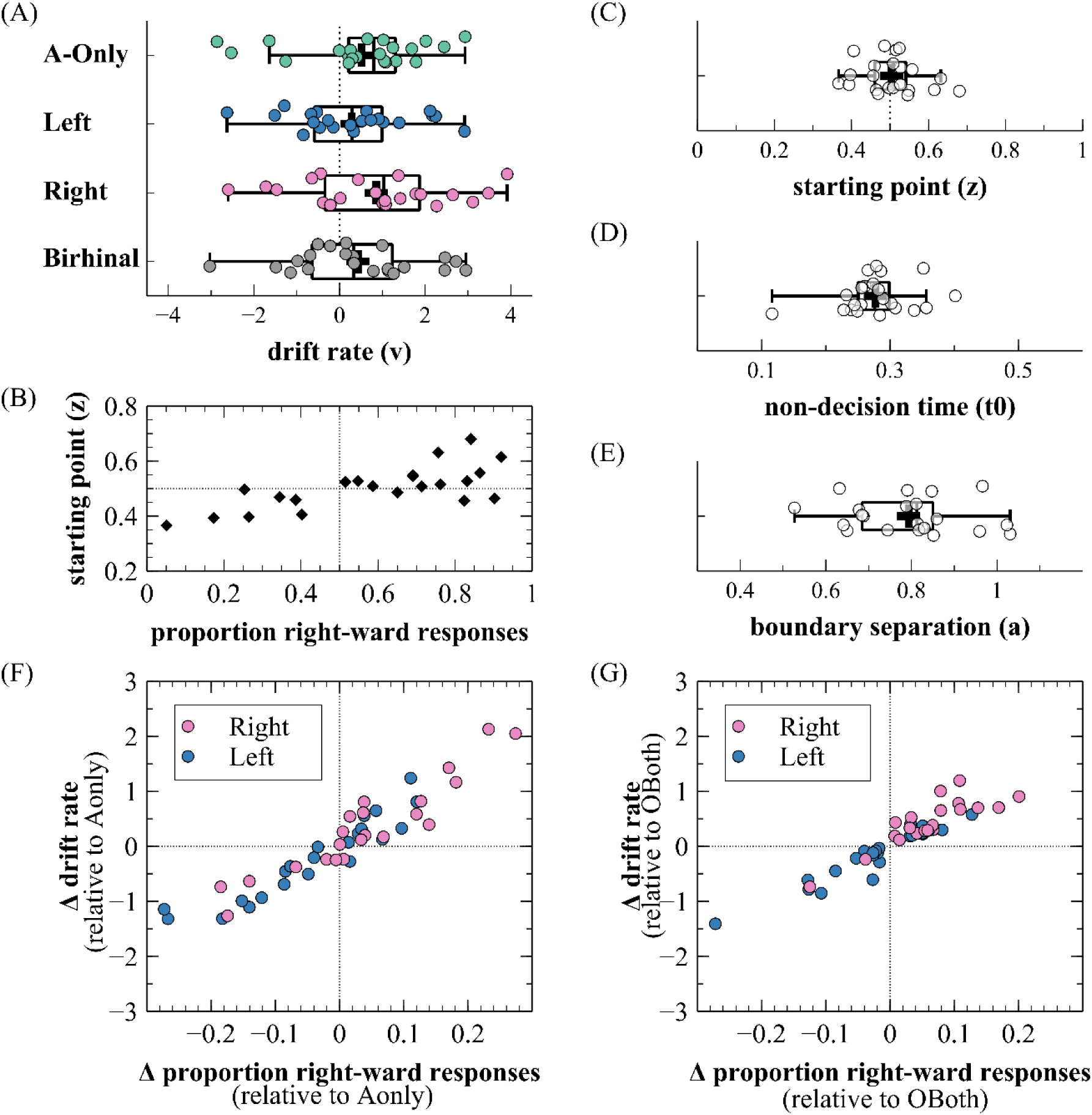
Diffusion Model Estimates for the Drift Model, allowing only drift rate to vary between conditions. Panel (A) depicts boxplots showing the distribution of drift rate estimates across conditions. The central line represents the median, the box spans the interquartile range (IQR), and whiskers extend to 1.5 times the IQR. Plus symbols denote the condition mean. Panel (B) illustrates the relationship between starting point (z) estimates and the proportion of right-ward responses. Participants with an overall bias towards the left response option (proportion right < 0.5) tend to have a starting point shifted towards the lower response boundary (i.e., z < 0.5), whereas participants with an overall bias towards the right response option (proportion right > 0.5) tend to have a starting point shifted towards the upper response boundary (i.e., z > 0.5), illustrating that the model generally captures the pattern of behavioral responses. Panels (C), (D), and (E) depicts boxplots showing the distribution of starting point (z), non-decision time(t0) and boundary separation (a), all of which were not allowed to vary between conditions. Panels (F) and (G) illustrate the relationship between the relative differences in drift rate and response proportions for Odor-Left and Odor-Right trials relative to controls.

### S5. Supplementary Analysis of Alpha Lateralization

**Table S8.** One-sided pair-wise comparisons contrasting ALI between monorhinal (congruent vs. incongruent) stimulation and controls (a-only, birhinal) at intermediate (± 5° azimuth) and outer (± 15° azimuth) sound positions.

|  | T | DF | P | P<sub>CORR</sub> | D | 95% CI |
| --- | --- | --- | --- | --- | --- | --- |
| Congruent vs. Birhinal, outer | -0.810 | 24 | 0.213 | 0.532 | -0.162 | $[-\infty -0.171]$ |
| Congruent vs. A-only, outer | -0.573 | 24 | 0.286 | 0.572 | -0.115 | $[-\infty -0.217]$ |
| Incongruent vs. Birhinal, outer | 2.412 | 24 | 0.012 | 0.040 | 0.482 | $[0.130 \infty]$ |
| Incongruent vs. A-only, outer | 2.917 | 24 | 0.004 | 0.019 | 0.583 | $[0.221 \infty]$ |
| Congruent vs. Incongruent, outer | -3.083 | 24 | 0.003 | 0.019 | -0.617 | $[-\infty -0.251]$ |
| Congruent vs. Birhinal, intermediate | -0.299 | 24 | 0.384 | 0.640 | -0.060 | $[-\infty 0.270]$ |
| Congruent vs. A-only, intermediate | -0.113 | 24 | 0.456 | 0.651 | -0.023 | $[-\infty 0.307]$ |
| Incongruent vs. Birhinal, intermediate | -0.770 | 24 | 0.776 | 0.776 | -0.154 | $[-0.483 \infty]$ |
| Incongruent vs. A-only, intermediate | -0.759 | 24 | 0.772 | 0.776 | -0.152 | $[-0.481 \infty]$ |
| Congruent vs. Incongruent, intermediate | 0.685 | 24 | 0.750 | 0.776 | 0.137 | $[-\infty 0.466]$ |

**Table S9.** Alpha Lateralization Index (ALI): 4 × 2 × 2 rmANOVA including the factors *condition*, *sound position*, and *sequence half*.

|  | DF | F | P | N<sub>p</sub><sup>2</sup> |
| --- | --- | --- | --- | --- |
| Condition | [3, 72] | 1.0478 | .377 | 0.042 |
| Sound position | [1, 24] | 11.622 | .002** | 0.326 |
| Sequence half | [1, 24] | 0.002 | .963 | < 0.001 |
| **Condition x sound position** | **[3,72]** | **2.904** | **.041\*** | **0.108** |
| Condition x sequence half | [3,72] | 0.121 | .947 | 0.005 |
| Sound position x sequence half | [1,24] | 5.218 | .031* | 0.179 |
| Condition x sound position x sequence half | [3,72] | 1.323 | .274 | 0.052 |

